# Psychometric Validation of the Education and Assessment of Genetic Literacy (EAGL) Measure

**DOI:** 10.64898/2026.05.22.727229

**Authors:** Lily S. Barna, Yi Liao, Michael R. Wierzbicki, Gabriela M. Ramírez-Renta, Kimberly A. Kaphingst, Chris Gunter

## Abstract

Genetic literacy is an integral measure for examining society’s interaction with genetics, but widely-used “genetic literacy” measures lack both knowledge comprehension measures and psychometric validation. To address these issues, we validated the Education and Assessment of Genetic Literacy measure (EAGL) in a sample of 2708 US participants, using both exploratory and confirmatory factor analysis. In addition to standard subjective and objective knowledge subscales, our measure’s distinct knowledge comprehension subscale focuses on autism as an example of a complex condition. Regression analyses showed a statistically significant interaction when looking at education and personal connection to autism in relation to knowledge comprehension (F=3.68, p=0.003). Separately, those in our sample with a connection to autism scored higher on the subjective knowledge section (F=19.52, p<0.001) only, concurring with previous demonstrations of a subjective-objective knowledge gap in science literacy. We explored geographic location as one potential factor in genetic literacy and found that metropolitan vs non-metropolitan status had no significant main effects on overall levels. After the validation process, we have two multi-domain measures which accurately capture the construct of genetic literacy and are available for wide use: the multi-faceted EAGL-long, which has previously been tested in thousands of participants, or the validated three-factor EAGL-short.

## INTRODUCTION

As accessibility to individual genomic information through genetic testing increases and raises corresponding healthcare implications, there is an equivalent need to increase genetic literacy. Genetic literacy, defined as “the sufficient knowledge and understanding of genetic principles for individuals to make decisions that sustain personal well-being and effective participation in social decisions on genetic issues,” is an integral measure to examine in the population and potentially improve in those receiving genetic testing.^1,2^ Given its importance, it is crucial to have a validated, effective, and reliable measure to measure genetic literacy levels.

Genetic literacy exists in a larger expanse of literacies and understandings that may interconnect in a variety of ways. Health literacy, defined as “the ability of an individual to obtain and translate knowledge and information in order to maintain and improve health in a way that is appropriate to the individual and system contexts,” has been researched individually and in connection with genetic literacy.^3^ Numeracy, defined as “how facile people are with basic probability and mathematical concepts,” has been widely associated with how people perceive health risks, linking it with health literacy.^4^ Research shows those with lower health literacy and numeracy may struggle to process genetic information in both print and oral forms, yet genetic literacy still diverges from other forms of literacy as a distinct but interrelated concept.^5,6^ Other studies have shown a lack of significant relationships between health literacy and genetic literacy using discriminant validity, or the ability of a measure to distinguish between differing constructs, to confirm genetic literacy to be distinct from both health literacy and numeracy.^7,8^

One difficulty for the field is that instruments attempting to capture genetic literacy vary widely, in fact capturing different subsets of genetic knowledge and literacy. Validated instruments such as the Rapid Estimate of Adult Literacy in Genetics (REAL-G) focus on familiarity with genetic terms, assessed through pronunciation.^9^ The Genetic Literacy and Comprehension measure (GLAC) built on the REAL-G to assess familiarity, instead functioning via 7-point Likert scale wherein participants rate their familiarity with commonly found genetic terms. Additionally, participants utilize Cloze technique multiple-choice questions to assess word comprehension.^10^ The validated Genetic Knowledge Index (GKI) focuses solely on objective knowledge, asking true/false questions rooted in core genetic conceptual understanding.^11^ Measures such as Ishiyama et al. ^12^ investigate knowledge of genetic terminology (measured via familiarity, or subjective knowledge); factual or objective understanding of genetic concepts; and awareness of the benefits and risks present in regard to genomic studies. While the addition of benefit/risk analysis is unique, the focus on subjective and objective knowledge is a core component of many genetic literacy measures. Many genetic literacy measures borrow from, update, and amend preexisting instruments such as Fitzgerald-Butt et al. ^13^, updating and psychometrically evaluating the widely used genetic knowledge measure created by Jallinoja and Aro ^14^. Finally, the International Genetics Literacy and Attitudes Survey (iGLAS-GK) measures genetic knowledge, heritability estimates, and attitudes toward various aspects of genetics use in education, in relation to the environment, in disease treatment, and more.^15^

A systematic review and survey to examine correlations and differences amongst six of the most commonly encountered instruments by Daly and Kaphingst^7^ found critical gaps in current genetic literacy measures. The results indicated two severe deficiencies in current measures: first, the lack of knowledge comprehension and applied knowledge among genetic literacy measures, and second, the lack of psychometric evaluation and validation amongst genetic literacy measures. Of the 89 studies examined, only two articles examined comprehension as an aspect of genetic literacy. Those two reports use the Genetic Literacy Survey (GLS^2^) composed by the Social and Behavioral Research Branch (SBRB) of the National Human Genome Research Institute (NHGRI)^2,16^, and later adapted by our unit into the Genetics and Autism Literacy Survey (GALS^1^). We then used GALS to examine genetic literacy both in the general population and a large autism genetics research study, noting that general population levels in the US have increased slightly in the decade since the GLS^1^. We also used GALS to examine other identity and belief factors contributing to genetic literacy levels; of those we examined, education level and confidence in one’s own genetic knowledge were the largest contributors.^17^

After the 2021 survey, we thoroughly reviewed our measure and ameliorated issues regarding language, content, and outdatedness of specific statements. We then added two more subscales to test other aspects of genetic literacy: applied knowledge and situational knowledge. While updated to reflect current language preferences and more accurate genetic understandings, the current Education and Assessment of Genetic Literacy measure (EAGL) still includes the key components of measuring genetic literacy from the original measure:^2^ familiarity (now called subjective knowledge), knowledge (now objective knowledge), and skills (now knowledge comprehension). Our subjective knowledge and applied knowledge sections come from the GLAC, utilizing both the Likert-scale ratings and multiple-choice Cloze technique questions.^10^ We swapped the term “abnormality” for “genome” in our survey to reflect our interests as the NHGRI, while shifting away from the normal/abnormal binary scholars have identified as problematic/ableist language.^18,19^ Our objective knowledge section derives from Jallinoja and Aro ^14^, with several changes made to account for updated research findings. Our knowledge comprehension section is rooted in our direct predecessor the GLS, with the significant change of shifting our infographic and comprehension questions from a focus on the *BRCA* gene family^2,16^ to the multi-factorial genetic and environmental contributors to an autism diagnosis. This version is available to researchers as the 46-item EAGL-long measure.

The lack of thorough psychometric validation represents a lack of clarity in overall genetic literacy measurement. Clarity is needed in many steps of measuring genetic literacy: from defining the term itself, to distinguishing between subjective and objective knowledge, and to capturing the critical component of knowledge comprehension. It is with this understanding of the gaps present in genetic literacy measures that this study aimed to validate the EAGL measure as a thorough and widely applicable instrument that accurately gauges genetic literacy levels amongst varying populations, and includes the crucial component of knowledge comprehension.

We conducted an iterative validation process in three phases collected from December 2024 through February 2025. Following data collection, we performed Exploratory Factor Analysis (EFA) and Confirmatory Factor Analysis (CFA) to refine the instrument, resulting in a validated, three-factor version: the EAGL-short. Our results demonstrate that the EAGL-short effectively captures three core constructs of genetic literacy: subjective knowledge, knowledge comprehension, and conceptual knowledge/objective knowledge. These findings provide researchers and clinicians a validated tool for measuring genetic literacy.

## MATERIALS AND METHODS

### Sample

All participants provided informed consent before participating in the survey and received compensation of $5.00. The survey was conducted entirely online via SurveyMonkey (to which participants were directed from the Prolific site), with an average completion time ranging from 10-20 minutes depending upon the additional ad-hoc questions included in the survey. The study was determined to be exempt human subjects research by the National Institutes of Health Institutional Review Board (IRB002137 / MOD008338).

We collected data in three sequential survey waves between July 2024 and February 2025, with all participants recruited via the online platform Prolific (see web resources). We utilized a sequential sampling strategy to maximize our statistical power and have samples for both Exploratory Factor Analysis (EFA) and Confirmatory Factor Analysis (CFA). The first two samples (n = 1005 and n = 702) were combined for the Exploratory Factor Analysis (EFA), while the third sample (n=1001) was reserved for Confirmatory Factor Analysis (CFA) to validate the factor structure identified in the EFA. Summary demographic details for the combined sample can be found in Table 1, with more complete details for each run in Table S1.

**Table 1.** Participant characteristics for combined sample (N = 2708)

| Table 1. Participant characteristics for combined sample (N = 2708) |  |  |  |
| --- | --- | --- | --- |
| Variable | Category | Frequency | % Valid |
| Age | 18-25 | 436 | 16.1% |
|  | 26-39 | 962 | 35.5% |
|  | 40-49 | 538 | 19.9% |
|  | 50-59 | 422 | 15.6% |
|  | 60-69 | 272 | 10.0% |
|  | 70+ | 78 | 2.9% |
| Education | Kindergarten through 8th grade | 5 | 0.2% |
|  | Some high school, no diploma | 41 | 1.5% |
|  | High school diploma or equivalent | 1,095 | 40.4% |
|  | Associate degree | 407 | 15.0% |
|  | Bachelor's degree | 785 | 29.0% |
|  | Professional degree | 311 | 11.5% |
|  | Doctorate degree | 64 | 2.4% |
| Connection to Autism | No | 2,136 | 78.9% |
|  | Yes | 572 | 21.1% |
| Metro Status | Metro | 2374 | 87.67% |
|  | Nonmetro | 330 | 12.19% |
|  | Invalid Responses | 4 | 0.15% |

Table 1 displays the demographic characteristics of the total study sample (N = 2708). Age groups range from 18 to 70+ years. Education levels span from elementary through doctoral education. Connection to Autism indicates whether participants reported any personal or familial connection to autism. Metro Status categorizes participants' geographic location as metropolitan or non-metropolitan based on Federal Information Processing System (FIPS) Codes for States and Counties and Rural-Urban Continuum Codes (RUCC, see web resources).

The first sample (EAGL1, n = 1005) was conducted in two phases, with 302 responses collected on July 24, 2024, and 721 responses collected between July 31 and August 4, 2024. This was done to establish the proficiency of Prolific and ensure we could obtain high-quality data and explore how long data collection would take. This sample was designed to be nationally representative of the adult, English-speaking population in the United States according to gender, age, and ethnicity. The nationally representative proportions are calculated from US Census Bureau population group estimates from 2021.^20^ After removal of incomplete and invalid responses, 1005 participants remained in the final dataset.

We found that 57.8% of the EAGL1 sample held a bachelor’s degree or higher education, which exceeded national representation. Given our own previous data demonstrating education as a moderating factor for genetic literacy overall,^17^ EAGL2 (n = 702) was collected on December 3-4, 2024, to oversample those with high school education and below.^21^ We introduced attention check questions into EAGL2 and 3 to help streamline the data cleaning process and ensure validity of responses. After removing incomplete and invalid responses from the 714 collected, along with those who failed the attention check question, we were left with 702 responses, with this sample containing 22.4% of participants with a bachelor’s degree or higher.

EAGL3 (n = 1001) was collected on February 19, 2025, and we designed it to match the combined previous two samples in educational distribution. A total of 1001 remained after we cleaned the original 1010 responses by removing incomplete and invalid responses, along with those who failed the attention check questions.

For regression analyses and other statistical tests, we utilized the combined dataset of all three samples to maximize statistical power. However, we had to remove four samples at this point because of invalid ZIP codes: one from EAGL1, two from EAGL2, and one from EAGL3; this produced N = 2704 for all tests after the EFA/CFA. We examined the consistency of scores across all knowledge domains to ensure appropriateness of combining the data. For confirmation, we ran a coefficient of variation (CV) analysis on the EAGL-long domains, which revealed excellent consistency across all knowledge domains, with CV values ranging from 0.29% for Subjective Knowledge to 1.35% for Knowledge Comprehension (Table S2). These exceedingly low values (all < 1.5%) indicate minimal variation across samples, supporting our choice to combine the datasets for analysis.^22^

### Measures

#### Genetic Literacy

We measured genetic literacy using the Evaluation and Assessing Genetic Literacy measure (EAGL), first developed by Abrams et al^2,16^ as the Genetic Literacy Scale (GLS) and later adapted by our laboratory as the Genetics and Autism Literacy Scale (GALS).^1^ Each iteration expanded the measure, including replacing *BRCA*-related material with an autism-related infographic adapted from Hoang et al.^23^ and making amendments for language-preference issues, outdated research, and other content issues identified through in-house review. The full EAGL-long measure contains five subscales: subjective knowledge, applied knowledge, situational knowledge, knowledge comprehension, and objective knowledge (shown in Table S3).

To assess subjective knowledge, or familiarity, participants rate their familiarity with eight common genetic terms: genetic, chromosome, susceptibility, mutation, variation, genome, heredity, and sporadic. Utilizing a seven-point Likert scale, answers ranged from not at all familiar (1) to completely familiar (7).

The applied knowledge (previously called “familiarity”) section consists of eight corresponding multiple-choice questions, utilizing the same genetic terms from the subjective knowledge section in a sentence. Participants must fill in multiple choice questions in a Cloze-style technique, with only one correct choice per question.

We created the situational knowledge questions to reflect potential situations in which one needs genetic understanding. Participants answer six multiple choice questions with subtopics including Mendelian inheritance, polygenic traits, genetic testing purposes, gene therapy, cloning, and sex-linked traits.

The knowledge comprehension (previously called “skills”) section contains a pedigree with cups and balls demonstrating gene-environment interactions that can result in reaching the threshold for an autism diagnosis (adapted from Hoang et al^23^), as a model of a complex condition. Participants then are asked six questions, scoring one point for every correct answer. The infographic is available for viewing at any point while answering the questions. Questions cover topics such as the purpose of genetic testing in autism, the impact of environmental and genetic factors, and the likelihood of autism diagnoses amongst family members.

The objective knowledge (previously called “knowledge”) section is composed of 17 True/False questions centered on multiple concepts within genetics. Participants answer “True” or “False” for each statement, with topics covering gene-environment interactions, heredity and condition carriers, mutations, and similar core genomic concepts. Correct responses earn one point, while incorrect responses receive zero points, resulting in a score range of 0-17.

The complete EAGL-long questionnaire with all items, response options, and correct answers is provided in Table S3.

#### Other measures

We measured objective numeracy using a three-item measure from Lipkus et al. ^4^ Participants answered three multiple-choice questions that center on basic probability, turning a percentage into a proportion, and identifying risk magnitudes when presented as proportions.

Because our knowledge comprehension infographic and questions focus on autism, we asked the participants “Are you or is anyone in your immediate family autistic?” The possible answers to “connection to autism” were yes or no.

Metropolitan or non-metropolitan distinctions were made via Federal Information Processing System (FIPS) Codes for States and Counties and Rural-Urban Continuum Codes (RUCC, see web resources). A RUCC of 1-3 delineates a metropolitan area, with 4-9 delineating a nonmetropolitan county. The specific numbers indicate the level of adjacency to a metropolitan area, considering population of county and proximity to metro counties. As stated above, four participants had to be removed from regressions and statistical analyses as they provided invalid 5-digit zip codes.

### Statistical Analysis

Using R Statistical version 4.3.3 and v4.4^24^ and SAS (version 9.4), we performed data preparation with readr^25^, dplyr^26^, and tidyverse^27^; and used reshape2^28^ for data restructuring. We utilized the car package^29^ and proc glm for analysis of variance (ANOVA), multcomp^30^ for pairwise comparisons, and jtools^31^ and proc glm/logistic for regressions. CFA path analysis was conducted with lavaan,^32^ qgraph,^33^ and MASS.^34^ Data were visualized with ggplot2^35^ and ggpubr,^36^ and the subsequent analyses and visualizations were exported with writexl^37^ and broom.^38^ Missing data were handled via case-wise deletion.

We combined our first two samples (n = 1707) for EFA using Mplus 8 statistical software. For factual questions encompassing applied knowledge, situational knowledge, and objective knowledge items, we calculated accuracy rates. Items with accuracy rates exceeding 90% were removed prior to EFA, as they lack sufficient variability to discriminate between participants and may not contribute meaningfully to identifying distinct factors. Accuracy rates for all items in the final three-factor solution are presented in Table 2, with complete accuracy rates for all EAGL-long items available in Supplementary materials. For EFA, we employed direct oblimin rotation, a method that permits correlations between items, and set a fixed number of factors. The selection of items was based on Howard’s^39^ “40-30-20” guideline, requiring items to demonstrate a minimum primary factor loading of 0.40 while avoiding cross-loadings on other factors that exceed .30, except when the primary loading surpasses the secondary loading by at least 0.20.

**Table 2.**
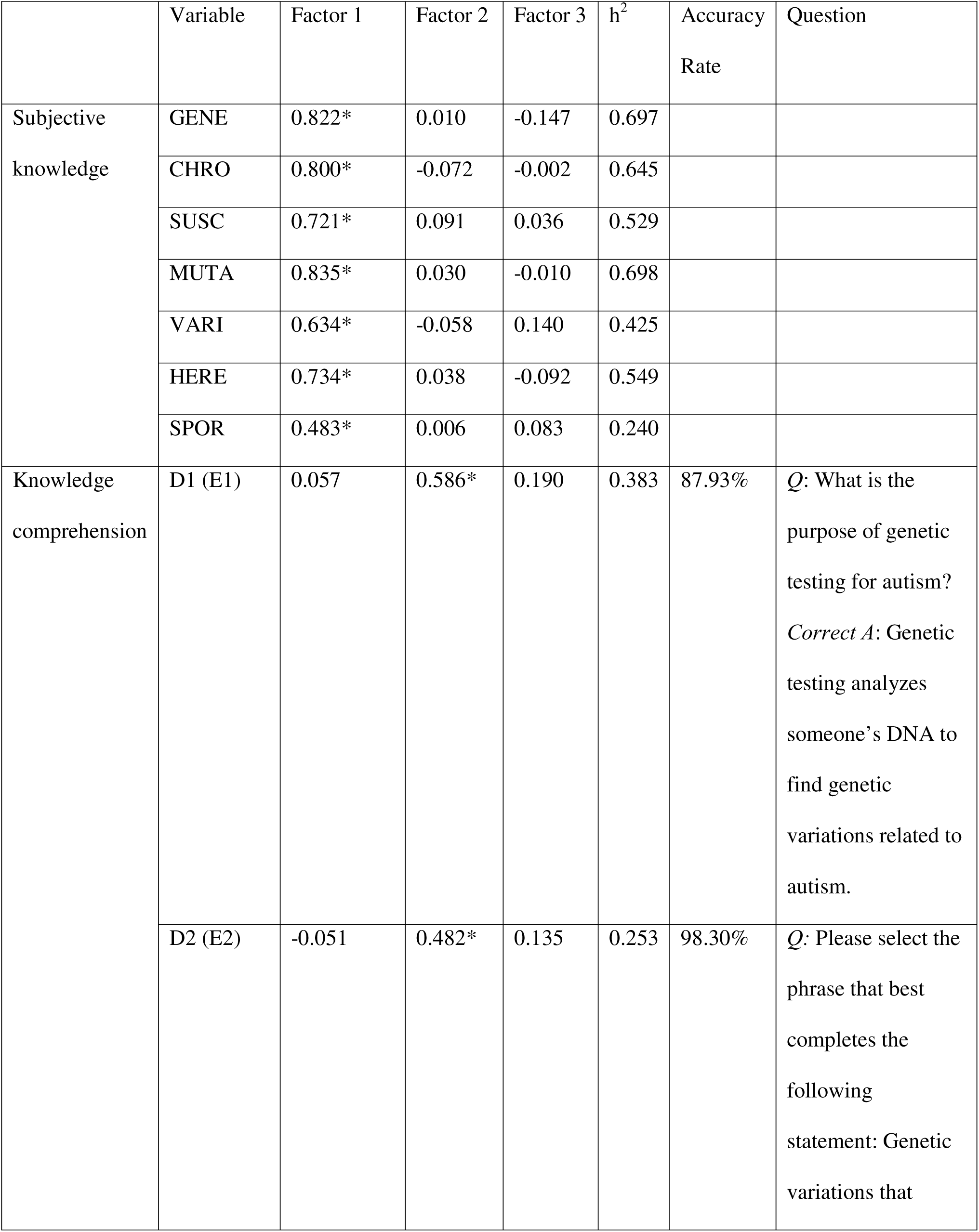
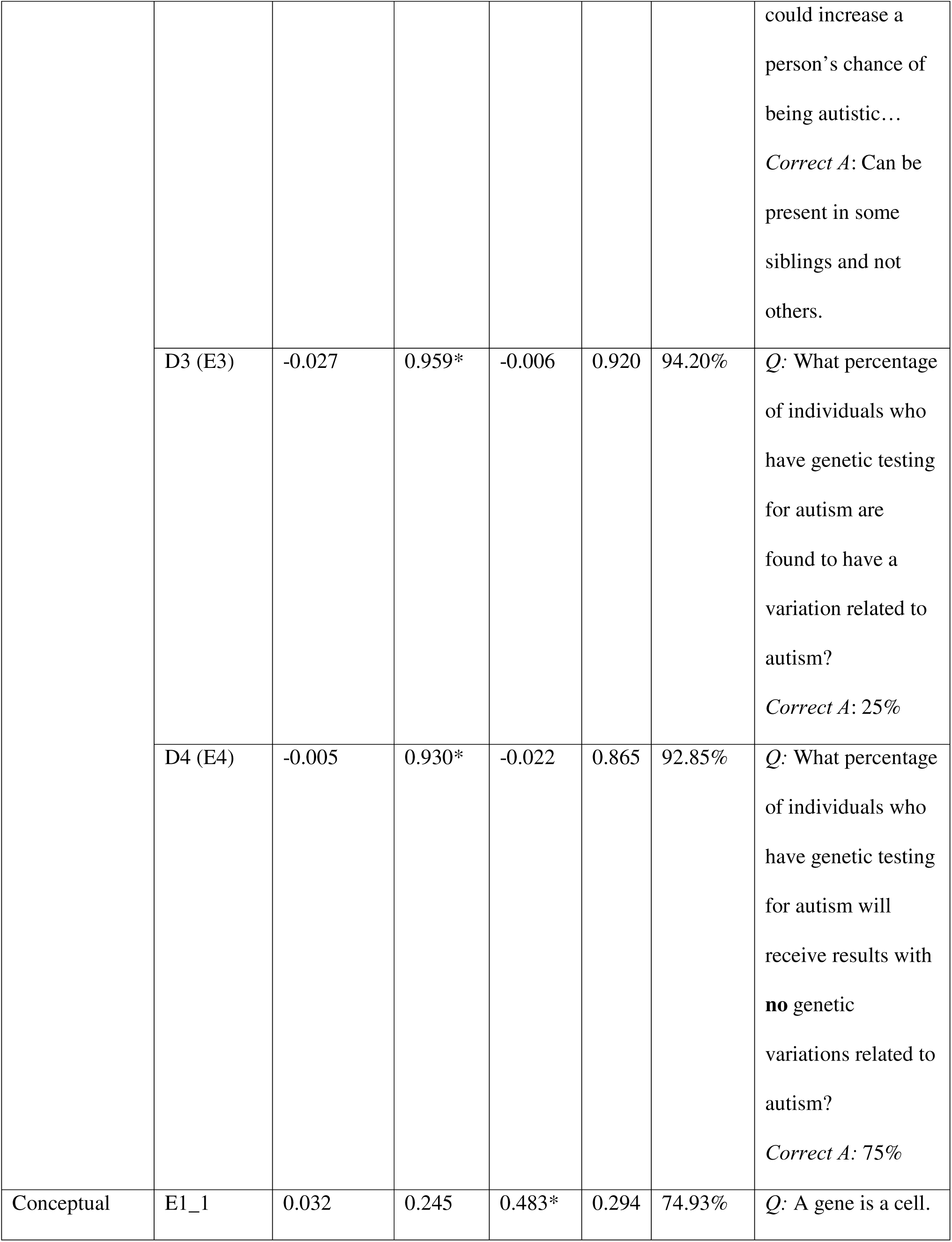
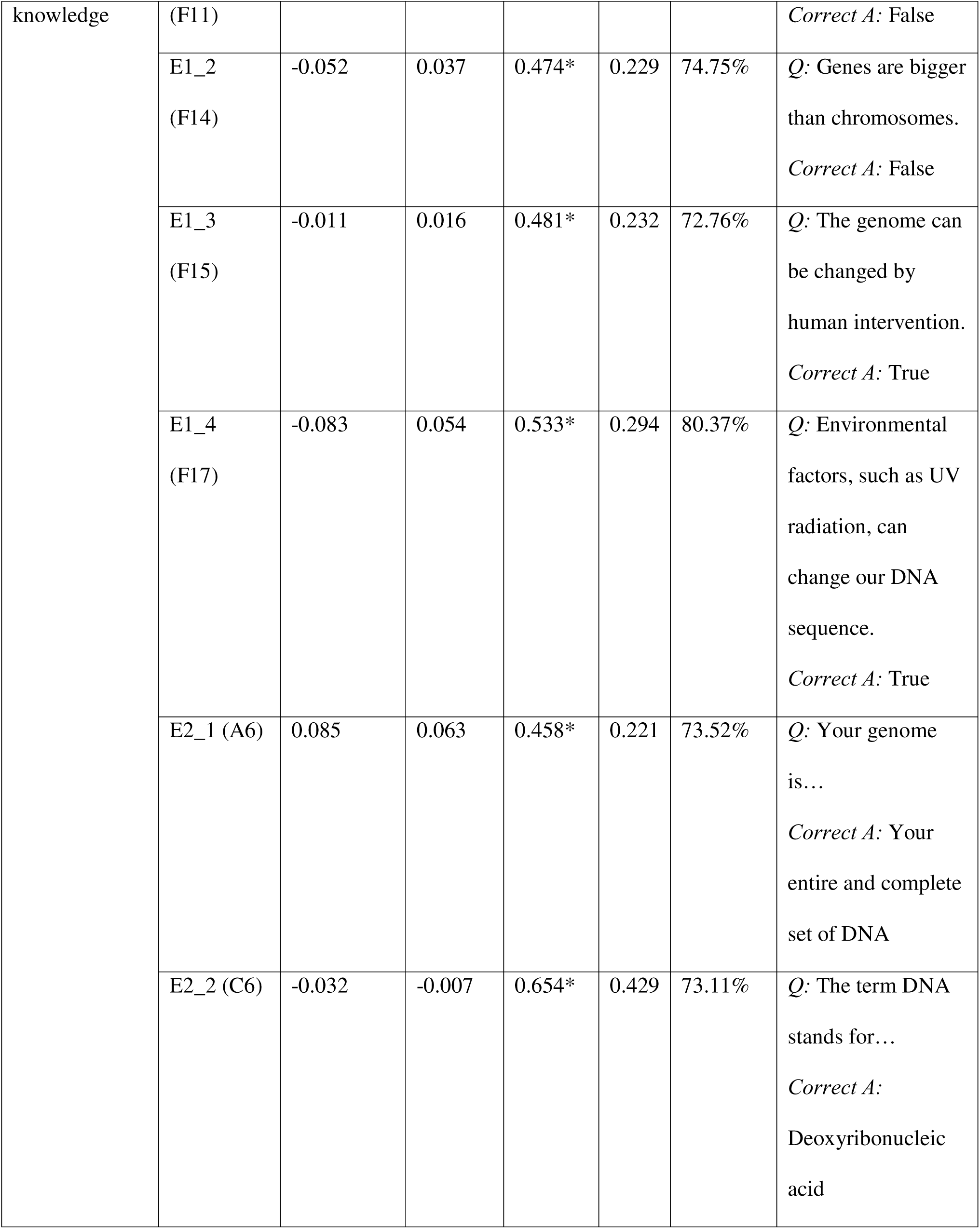
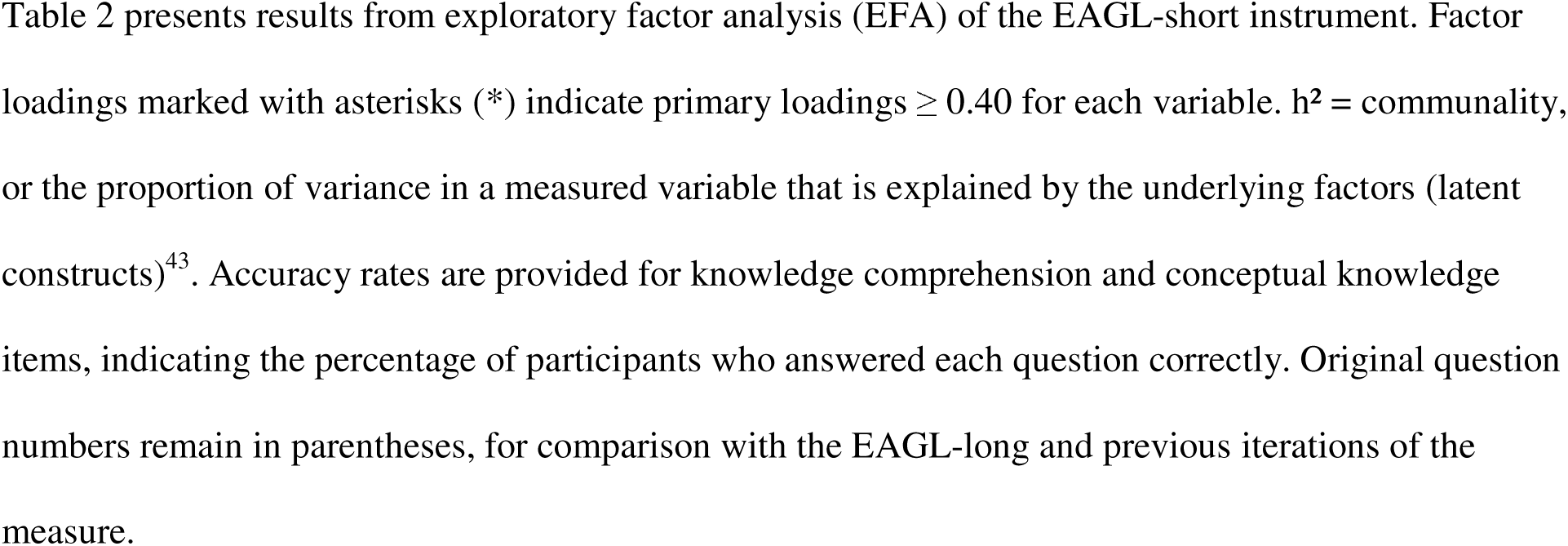
Factor Loadings, Communalities, and Accuracy Rates for EAGL-short (EFA)

We employed CFA to verify the factor structure discovered through EFA using the third sample (n = 1001). Due to the binary nature of some items, the weighted least squares mean and variance adjusted (WLSMV) estimator was used. This method assumes that observed categorical responses reflect an underlying continuous normal distribution segmented by threshold values that govern the transformation of continuous scores into categorical outcomes, as opposed to treating the categorical responses as inherently normally distributed.^40^ We used multiple indices to assess model fit: Comparative Fit Index (CFI), Root Mean Square Error of Approximation (RMSEA), and Standardized Root Mean Square Residual (SMRM). A CFI > 0.95, a RMSEA < 0.06, and a SRMR value of < 0.08 all indicate good model fit.^41^ Cronbach’s alpha was utilized to measure the internal consistency for each subscale, with a threshold of 0.70 or higher.

We chose EFA and CFA, as opposed to Item Response Theory (IRT), for several methodological reasons. Given that genetic literacy as a construct with multiple dimensions, factor analysis is optimal to account for those qualities. Additionally, EFA and CFA are more appropriate for measures with varying question types, as our measure includes Likert scale, true/false, numeric answer, and multiple-choice questions. The larger sample size allowed us to explore the relationships between variables we observed, as opposed to IRT which is more probabilistic. Lastly, CFA and EFA align with our theoretical framework, in which genetic literacy is comprised of distinct factors that interrelated with one another but also stand alone.

We performed linear regressions to examine associations between our genetic literacy outcome measures and predictor variables (age, education, connection to autism, numeracy, and metro status). Analysis of variance (ANOVA) was used to test the significance of effects and interactions in the regression models. Models included interaction terms between education and the other predictor variables, to see if educational effects varied amongst groups. Sensitivity analyses were performed to assess the robustness of the results of the ANOVA models. Finally, we performed cumulative logit regression models with the same predictor values and Kruskal-Wallis tests by each predictor.

For the categorical variable of education, the variable “Less than high school” serves as a reference level, with that group composed of all responses of “No schooling completed,” “Kindergarten through 8^th^ grade,” and “Some high school, no diploma” (n = 46). We performed education and metro status analyses on the full dataset (N = 2704), assessing model fit using F-statistics, p-values, and R² values.

## RESULTS

### Descriptive Statistics

The final combined sample (N = 2704) included participants across age groups, educational attainment groups, and metropolitan/nonmetropolitan groups (Table 1 for combined summary and Table S1 for more details). The largest age representation (35.5%) is in the 26-39 age bracket, while the largest educational grouping was those with a high school diploma or equivalent, 40.4% of the sample. 78.9% of participants reported no connection to autism either personally or in their family, while 21.1% reported having a connection. The majority (87.7%) of our participants resided in metropolitan areas. For connection to autism, 21.1% of our combined sample answered yes.

### Exploratory Factor Analysis (EFA)

We conducted EFA on the 46-item Education and Assessment Genetic Literacy measure (Table S3), initially identifying nine factors centering on various subject and knowledge domains (Table S4, Figure S1d). We ran EFA analyses fixing the factor number at five (Table S5), four (Table S6), and three (Table 2), and followed the “40-30-20” guideline for item selection. Heatmap representations of these four EFA analyses are in Figure S1 (panels a-d). We found three strong factors present throughout the measure, distilling the measure down to a three-factor solution with clear groupings: subjective knowledge, knowledge comprehension, and objective knowledge (Table 2). Items that did not load significantly onto any of the three factors (with loadings below 0.4) were identified for potential removal in the subsequent CFA phase, with the final validated survey becoming the EAGL-short. Eigenvalues and variance explained for all factors are presented in Table S6. We established a threshold of 0.4 as a baseline, with two items having loadings just under 0.4 but still retained due to the larger sample size and theoretical and practical relevance of the items.^42^

### Confirmatory Factor Analysis (CFA)

We performed CFA on the three-factor model identified through EFA, after removing items that failed to load significantly on any factor. We removed items A1-A5, A7, A8, C1-C5, E5, E6, F1-F10, F12, F13, F16, as well as the term “Genome,” as each item either loaded on multiple factors or failed to load on any factor with statistical significance. The resulting CFA model consists of 17 genetic literacy questions, constituting the EAGL-short (Figure 1). Factor loadings, standard errors, and standardized loadings for all items are presented in Table 3.

**Figure 1.**
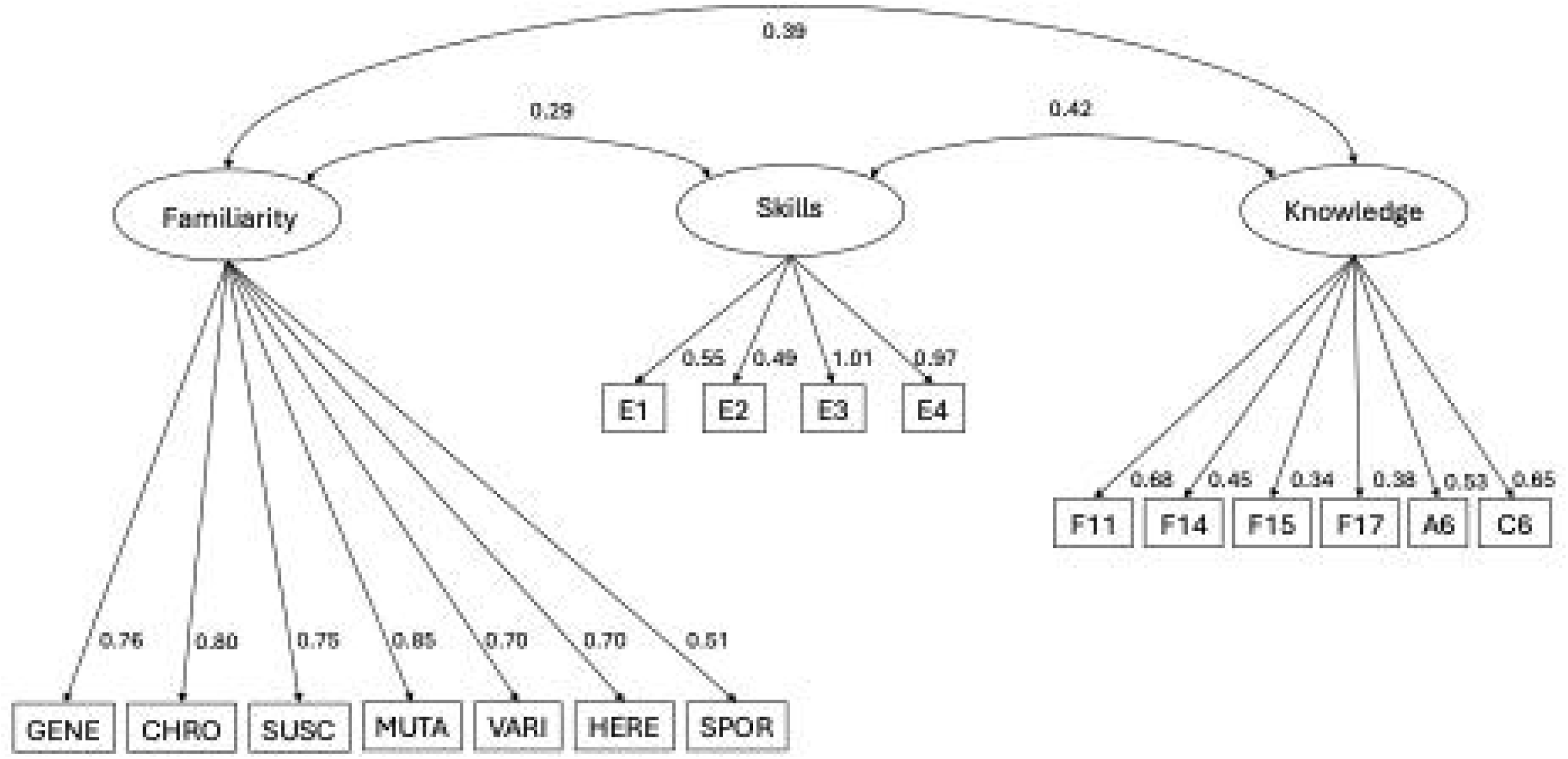
Confirmatory Factor Analysis path diagram for the EAGL-short, showing standardized factor loadings and correlations between factors. Model fit indices: CFI = 0.996, RMSEA = 0.031 [0.025, 0.037], SRMR = 0.080.

**Table 3.**
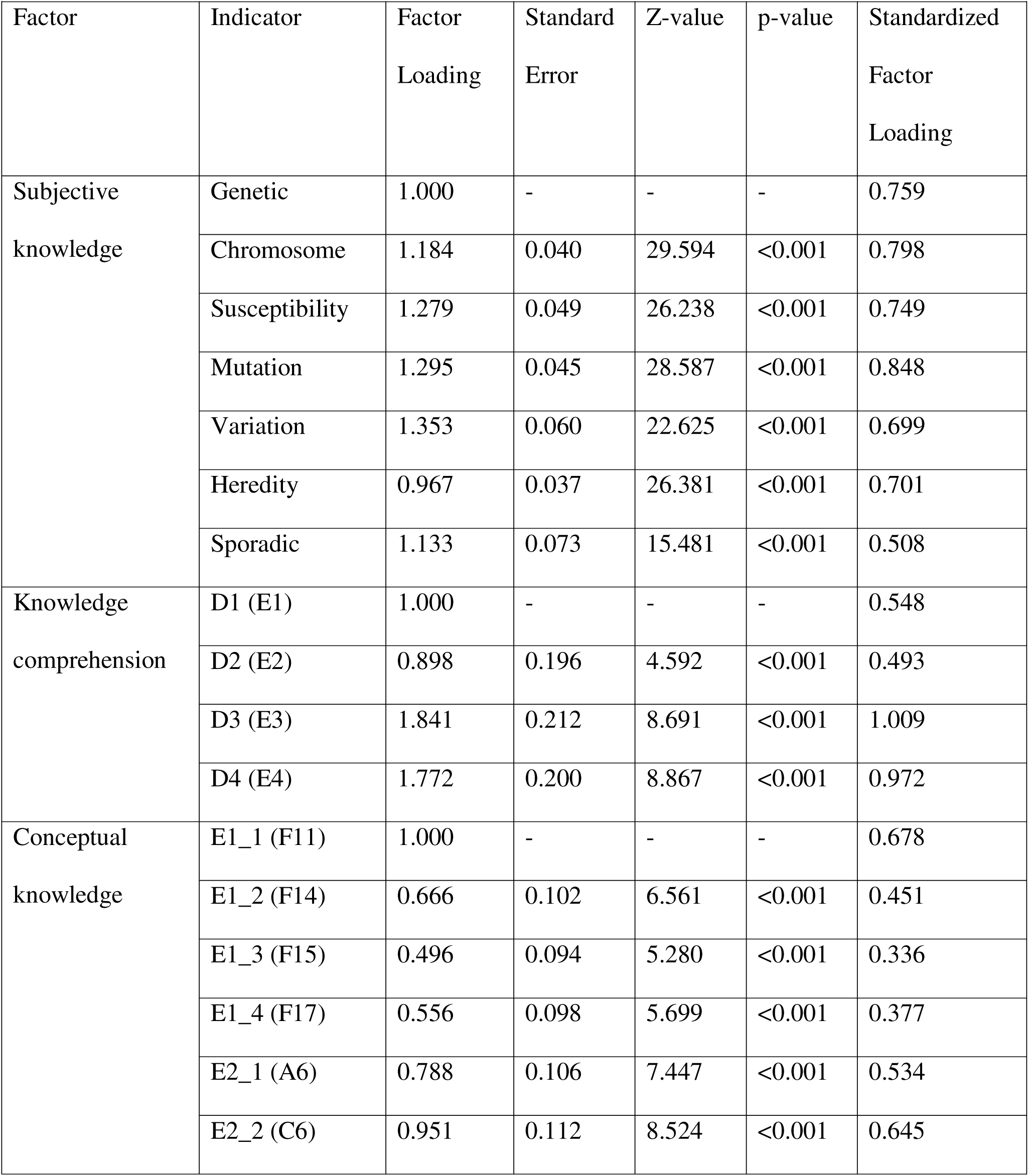
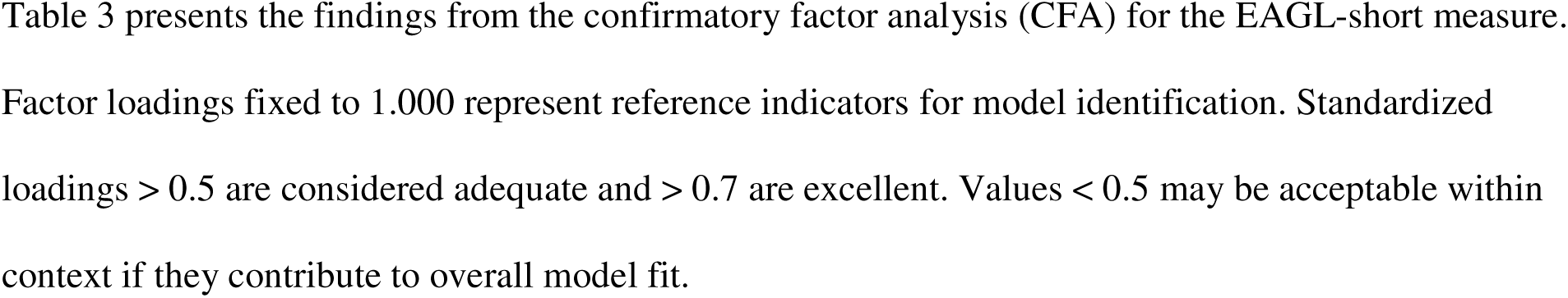
Factor loadings for EAGL-short (Confirmatory Factor Analysis)

The model demonstrated excellent fit across multiple indices. The CFI value (0.996 standard, 0.989 scaled) was above 0.95, indicating excellent fit. The RMSEA of 0.031 [0.025, 0.037] was below the 0.05 threshold, also suggesting good fit. The SRMR (0.080) is acceptable at the base threshold of ≤ 0.08. The chi-square test of model fit (*X*² (116) = 115.426, p = 0.498) was non-significant, indicating no significant differences between the model data and observed data, suggestive of an exceptional fit.

### Validated Measure

After our factor analysis, the EAGL-short demonstrated strong psychometric properties among three validated factors: subjective knowledge, knowledge comprehension, and conceptual knowledge. Subjective knowledge (Table S8, 7-item Likert scale; M = 5.69, SD = 0.95, α = 0.87), knowledge comprehension (Table S9, 4-item multiple-choice and numeric answer; M = 3.68, SD = 0.97, α = 0.83), and conceptual knowledge (Table S9, 6-item multiple-choice and true/false; M = 4.54, SD = 1.40, α = 0.67) all showed acceptable to excellent internal consistency (see Table S8 for detailed descriptive statistics of subjective knowledge items and Table S9 for frequency distributions of knowledge comprehension and conceptual knowledge items). Conceptual knowledge is an amalgamation of several questions from the applied knowledge, situational knowledge, and objective knowledge sections (A6, C6, F11, F14, F15, F17). The shared factor of conceptual knowledge appears to center on fundamental genetics principles of heredity and genetic changes that do or do not lead to disease. A potential reason for the relatively low α-value of 0.67 is the heterogeneity of the question types in this section, with True/False, multiple-choice, and multiple-choice fill-in-the-blank style questions.^44^

### EAGL-Long Performance

The mean values for all five portions of the EAGL-long are presented in Table S10. Compared to previous administrations of our genetic literacy survey, the results were slightly higher in this population.^1,17^ The EAGL-long subscales showed variable internal consistency, with subjective knowledge demonstrating good reliability (α = 0.877), while other subscales showed lower alpha values due to the heterogeneous nature of the items and smaller number of questions per subscale. Consistency across samples was confirmed through coefficient of variation analysis (Table S2), with all CV values below 1.5%.

### Regression Analysis

To examine how demographic variables predict the three constructs of genetic literacy, we performed regression analysis on the EAGL sample in its entirety, as well as on each of the three EAGL runs. The analysis centered on the EAGL-short subscales of subjective knowledge, knowledge comprehension, and conceptual knowledge.

Analysis of variance revealed several statistically significant associations between demographic variables and EAGL-short subscales (Table 4). Complete adjusted means for each demographic group are available in Table S11. Total numeracy score showed highly significant effects across all three subscales of EAGL-short (all p<0.001), consistently demonstrating the strongest associations. In contrast, metropolitan (RUCC 1-3) vs non-metropolitan (RUCC 4-9) status had no significant main effects on overall levels.

**Table 4.**
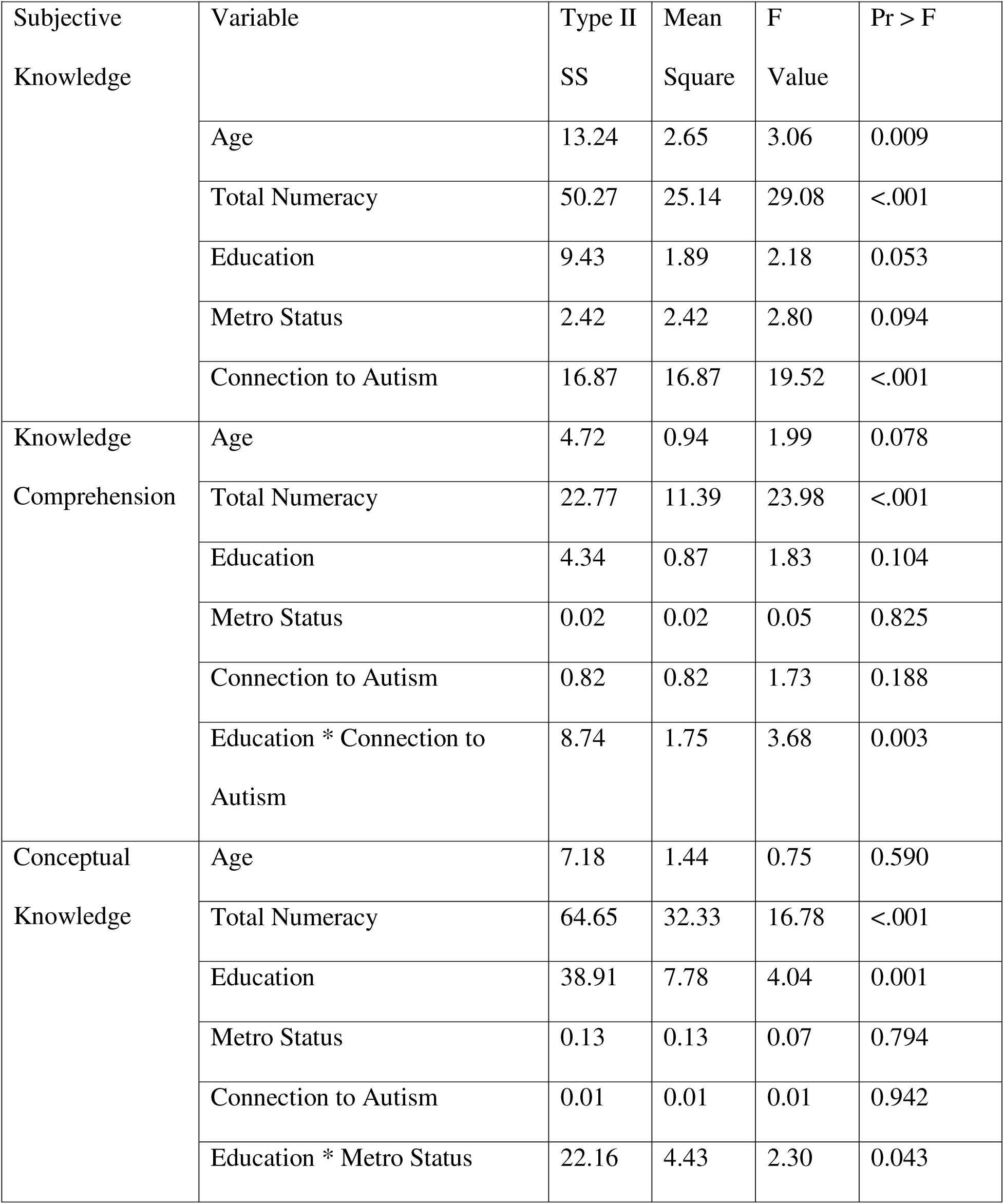
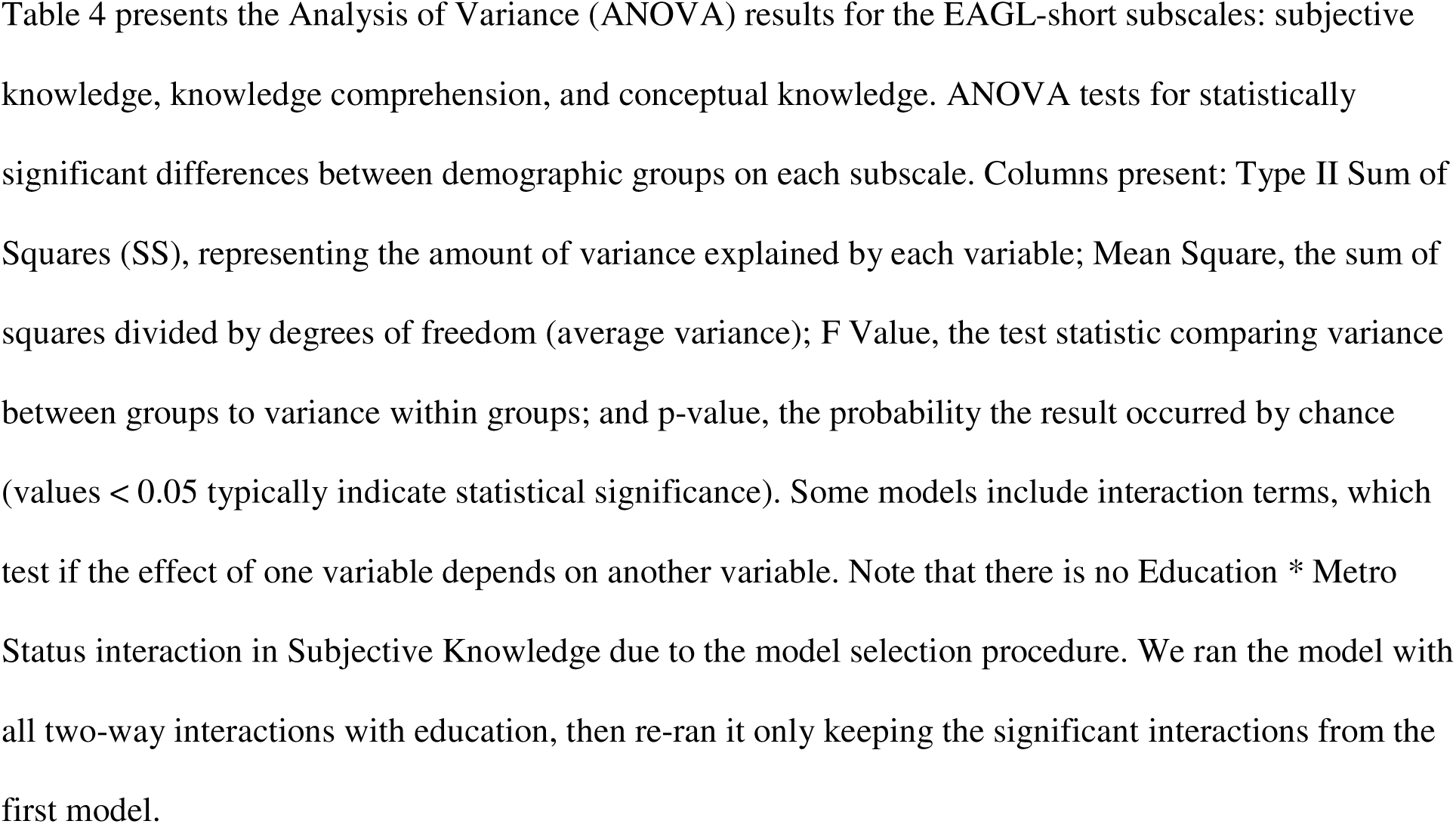
ANOVA Statistics.

For subjective knowledge (scored 0-7), age (F=3.06, p=0.009) and connection to autism (F=19.52, p<.001) also showed significant effects. Those with a connection to autism scored higher (adjusted M=5.97, Table S11) than those without (M=5.78), though the effect size was small (Cohen’s d=0.19), indicating limited practical significance. Connection to autism is determined by a question on the survey asking: “Are you or is anyone in your immediate family autistic?” This pattern, where those with a connection to autism showed higher subjective knowledge but not on other subscales, concurs with previous demonstrations of a subjective-objective knowledge gap in science literacy.^45^

For knowledge comprehension (scored 0-4), numeracy was the primary significant predictor (F=23.98, p<.001). Connection to autism showed no significant effect on knowledge comprehension (F=1.73, p=0.188, Cohen’s d=0.04). We did observe a statistically significant interaction between education and connection to autism (F=3.68, p=0.003).

For conceptual knowledge (scored 0-6), both numeracy (F=16.78, p<0.001) and education (F=4.04, p=0.001) showed significant main effects. Those with bachelor’s degrees (adjusted M=4.66) and doctoral degrees (M=4.63) scored notably higher than those with less than high school education (M=3.78). A statistically significant interaction between education and metro status was also observed (F=2.30, p=0.043).

## DISCUSSION

This study reports the psychometric evaluation and validation of the Education and Assessment Genetic Literacy measure (EAGL) in three robust U.S. population samples. The analyses indicate the EAGL-short instrument is sound and captures unique aspects of genetic literacy, a multidimensional construct that by definition should include a measurement of comprehension as opposed to only subjective or objective knowledge.

The exploratory and confirmatory factor analyses ultimately illustrate a highly factorial construct in the EAGL-long that touches on many dimensions of genetic literacy and education. The longer version is still composed of individually validated components and may serve in a variety of settings including academic pre- and post-tests or genetic testing population surveys, while the shorter version may be more streamlined for clinical or exit-survey settings. The psychometrically validated three-factor structure in EAGL-short, which highlights subjective knowledge, knowledge comprehension, and conceptual knowledge, aligns with the previous dimensions identified in both Little et al.^1^ and Abrams et al.^2^ The psychometric validation process identified several items amongst the five subscales in EAGL-long that either loaded on unique factor domains, or did not discriminately load on any factor domain, thereby not measuring any specific aspect of genetic literacy. The EAGL-short condensed the more varied EAGL-long into a tight-knit body of effective questions.

Findings from a validation study of the EAGL predecessor showcase via discriminant validity analysis that genetic literacy is a distinct construct, separate from both health literacy and numeracy.^8^ While it is clear these constructs coexist and likely relate to one another, genetic literacy uniquely hones in on the processing of genetic health information. Liao et al.^8^ emphasize the need to examine these constructs both in connection with one another and separately to glean their truest effects on health decision-making.

The knowledge comprehension factor represents an integral and underexplored component of genetic literacy. With many previous measures focusing solely on subjective and/or objective knowledge, the EAGL directly measures individuals’ ability to synthesize and apply given genetic information. The strong scores throughout the section indicate that with cohesive and clear communication, synthesizing genetic information can be an equitable and universal accomplishment. Specifically using an example of the complex-condition model in autism showcases how individuals can understand complex scientific ideas, such as gene-environment interactions, effectively. Further research should investigate the longevity of their understanding of the information they synthesized in the knowledge comprehension section.

The overall scores in this study were higher than those of the previous measure iterations.^1,17^ This may indicate higher genetic literacy levels overall consistent with Little et al.^1^ finding that genetic literacy levels are rising amongst the general population over time, with continued room for improvement. It may also reflect increased public exposure to genetic information, as our previous data collection occurred in 2021, and genomics has only further made it to the forefront of mass media. Or, it could reflect qualities of the users of the Prolific platform compared to our previous public panels.

The findings that those with a personal or familial connection to autism scored higher on subjective knowledge, but not knowledge comprehension or conceptual knowledge, provides evidence for the subjective-objective knowledge gap present in health literacy broadly. The subjective-objective knowledge gap posits that the more people believe they understand the science, the more confident they may feel in their understanding of it.^45,46^ Those with connections to autism may hold stronger feelings toward genetics, be it positive or negative, thereby finding themselves more familiar with genetic terms, even if they do not actually know other aspects of genetics.^45^ Another potential reason for higher subjective knowledge is more exposure to genetic terms in their daily lives. Interactions with genetic counselors and genetic testing, increasingly common components of receiving an autism diagnosis, may contribute to increased familiarity with genetic terms.

The statistically significant interaction between education and connection to autism in relation to knowledge comprehension suggests that one’s educational attainment may moderate how their personal connections relate to their genetic literacy levels. Those with higher educational attainment may have additional skills to interpret/understand genetic information that they have encountered through their personal or familial autism connection, whereas those with less education may have less skills through which their personal experiences can translate into higher genetic literacy. Another potential explanation is that those with higher educational levels may seek out more genetic information related to their autism connection and may have stronger comprehension skills overall regarding complex genetic information.

There were no main effects of metropolitan status on any of the three genetic literacy subscales, suggesting that geographic location alone does not predict genetic literacy levels. A statistically significant interaction between education and metro status was observed for conceptual knowledge (F=2.30, p=0.043), though the specific pattern of this interaction requires further investigation to interpret.

While we hypothesized that those in metropolitan areas would have higher genetic literacy due to greater exposure to genetic concepts through healthcare services, testing opportunities, genetic counselors, and research institutions, our data did not support this hypothesis. The lack of main effects suggests that deficits in genetic literacy may be more universal than geography-specific. However, the observed interaction may indicate that the relationship between education and conceptual knowledge differs between metropolitan and nonmetropolitan contexts, potentially reflecting differences in educational quality, curriculum emphasis, or access to genetics-related healthcare services. Further research should investigate both the nature of this interaction and potential differences in genetic training among healthcare providers across geographic areas.

Through these validation efforts, the EAGL is ready for widespread use in a variety of social populations and settings. Still, further research could help establish its full efficacy in multiple social groups and languages. There is additional room for streamlining, and reorganization of the survey questions may also be a potential avenue of exploration, helping to further explicate the mediation and moderation of subjective knowledge, knowledge comprehension, and conceptual knowledge on one another. The findings that genetic literacy concepts can be accessible across educational, age, and geographical groups, has important implications for provider education, patient education, mass communications, genetic counseling, public health initiatives, and other groups that interact with genetic information. For those interested in the original scale and domain structure, the full EAGL-long remains available for use as a comprehensive coverage of genetic literacy, though researchers should be aware of the psychometric limitations of individual subscales.

The validation of a genetic literacy measure that captures subjective knowledge, objective knowledge, and knowledge comprehension has implications for both the continued field of genetic literacy research and the public as genetics continues to grow in clinical utilization. Understanding individuals’ genetic literacy levels can inform providers about effective communication techniques and educational interventions. Psychometric validation allows researchers to accurately gauge the complex genetic literacy levels of any population they wish to survey, helping to create more targeted and productive genetic communication interventions and educational materials.

## Limitations

While this study provides robust validation of the EAGL measures, several limitations should be acknowledged. Utilizing an online survey recruitment platform may lead to selection bias and exclusion of those without access to technology.^47,48^ The self-reporting of subjective knowledge may lead to implicit biases, with some overestimating or underestimating their own familiarity levels.^49^ The use of autism as our example of the complex conditions model may introduce condition-specific biases, with those who have opinions on or prior experience with autism. Researchers can address these limitations in future studies by surveying in-person or over the phone to reach participants without internet capabilities; introducing a more objective measurement of term recognition; or selecting a different example to illustrate the complex condition.^2^ Additionally, this validation was run with only U.S. participant who speak English as their first language. Running the validation in additional languages and locations could help increase the spread of the measure, ensuring it correctly captures genetic literacy across places and languages.

## Conclusion

This study ultimately validates the Education and Assessment Genetic Literacy measure as a psychometrically sound measure of genetic literacy. The analyses showcase a robust and multi-factorial long form version that touches on many domains of genetic knowledge and literacy, and a streamlined shortened version that encompasses three key factors of genetic literacy: subjective knowledge, knowledge comprehension, and conceptual knowledge. These factors are recognized as core constructs of genetic literacy, further solidifying EAGL as a measure suitable for widespread use. Knowledge comprehension, or skills, has been identified as a unique and valid construct, something absent from many previous genetic literacy measures but worth utilizing in forward studies and interventions. With so much discrepancy amongst current genetic literacy definitions, measures, and implications, creating and identifying psychometrically sound instruments is of high importance.

While previous literature asserts a continued effect of education on genetic literacy levels, the results of this survey indicate an overall increase in score regardless of educational level, along with a nonsignificant difference in score amongst educational levels. The additional nonsignificant difference in score amongst those in metro and non-metro areas indicates that genetic literacy issues pervade educational and geographical strata. Genetic literacy needs to be evaluated and improved across the board to help empower patients and providers alike and increase informed decision making and having sound and robust surveys is a critical step in identifying current genetic literacy levels and gaps that need addressing.

## Data and code availability

The raw data utilized in this study were derived from a larger research project. The dataset will be publicly available after the initial articles reporting on the collected data are published. Summary statistics are published in Little et al. 2022^1^. Until then, the dataset can be accessed by contacting the corresponding author upon reasonable request.

## Acknowledgements

This project was funded by a SPARK Research Match grant (RM0149) and the National Institutes of Health: National Human Genome Research Institute Intramural Research Program (HG200410-01). The contributions of the NIH authors are considered Works of the United States Government. The findings and conclusions presented in this paper are those of the authors and do not necessarily reflect the views of the NIH or the U.S. Department of Health and Human Services. We thank the Gunter and Persky labs at NHGRI for suggestions throughout.

## Author contributions

Conceptualization and design of experiments: L.S.B., Y.K., G.M.R.R., K.A.K., C.G. Data analysis: L.S.B., Y.K., M.R.W., K.A.K., C.G. Data curation: L.S.B., Y.K., G.M.R.R., M.R.W. Writing-original draft: L.S.B., C.G. Writing-review & editing: L.S.B., Y.K., M.R.W., K.A.K., C.G. All authors have reviewed and approved the final version of the manuscript.

## Declaration of generative AI and AI-assisted technologies in the writing process

During the preparation of this work, the authors used Claude v4.1 Opus in order to check for grammar and generate visualizations. After using this tool/service, the authors reviewed and edited the content as needed and take full responsibility for the content of the publication.

## Supplemental information

Supplemental Information includes one file with additional data tables, figures, and analyses.

Web Resources section

Prolific https://www.prolific.com

USDA Rural-Urban Continuum Codes or RUCC https://www.ers.usda.gov/data-products/rural-urban-continuum-codes/documentation

## References

1. Little, I.D., Koehly, L.M., and Gunter, C. (2022). Understanding changes in genetic literacy over time and in genetic research participants. Am J Hum Genet 109, 2141–2151. 10.1016/j.ajhg.2022.11.005.

2. Abrams, L.R., McBride, C.M., Hooker, G.W., Cappella, J.N., and Koehly, L.M. (2015). The many facets of genetic literacy: Assessing the scalability of multiple measures for broad use in survey research. PLoS ONE 10, 1–11. 10.1371/journal.pone.0141532.

3. Liu, C., Wang, D., Liu, C., Jiang, J., Wang, X., Chen, H., Ju, X., and Zhang, X. (2020). What is the meaning of health literacy? A systematic review and qualitative synthesis. Fam Med Community Health 8. 10.1136/fmch-2020-000351.

4. Lipkus, I.M., Samsa, G., and Rimer, B.K. (2001). General Performance on a Numeracy Scale among Highly Educated Samples. Medical Decision Making 21, 37–44. 10.1177/0272989x0102100105.

5. Lea, D.H., Kaphingst, K.A., Bowen, D., Lipkus, I., and Hadley, D.W. (2011). Communicating genetic and genomic information: health literacy and numeracy considerations. Public Health Genomics 14, 279–289. 10.1159/000294191.

6. Kaphingst, K.A., Blanchard, M., Milam, L., Pokharel, M., Elrick, A., and Goodman, M.S. (2016). Relationships Between Health Literacy and Genomics-Related Knowledge, Self-Efficacy, Perceived Importance, and Communication in a Medically Underserved Population. J Health Commun 21 *Suppl 1*, 58–68. 10.1080/10810730.2016.1144661.

7. Daly, B.M., and Kaphingst, K.A. (2023). Variability in conceptualizations and measurement of genetic literacy. PEC Innov 2, 100147. 10.1016/j.pecinn.2023.100147.

8. Liao, Y., Wei, W., Barna, L.S., Gunter, C., and Kaphingst, K.A. (2025). Genetic literacy scale: Construct and discriminant validity and population differences. Patient Educ Couns 142, 109390. 10.1016/j.pec.2025.109390.

9. Erby, L.H., Roter, D., Larson, S., and Cho, J. (2008). The rapid estimate of adult literacy in genetics (REAL-G): a means to assess literacy deficits in the context of genetics. Am J Med Genet A 146a, 174–181. 10.1002/ajmg.a.32068.

10. Hooker, G.W., Peay, H., Erby, L., Bayless, T., Biesecker, B.B., and Roter, D.L. (2014). Genetic literacy and patient perceptions of IBD testing utility and disease control: a randomized vignette study of genetic testing. Inflamm Bowel Dis 20, 901–908. 10.1097/mib.0000000000000021.

11. Furr, L.A., and Kelly, S.E. (1999). The Genetic Knowledge Index: Developing a Standard Measure of Genetic Knowledge. Genetic Testing 3, 193–199. 10.1089/gte.1999.3.193.

12. Ishiyama, I., Nagai, A., Muto, K., Tamakoshi, A., Kokado, M., Mimura, K., Tanzawa, T., and Yamagata, Z. (2008). Relationship between public attitudes toward genomic studies related to medicine and their level of genomic literacy in Japan. American Journal of Medical Genetics Part A 146A, 1696–1706. 10.1002/ajmg.a.32322.

13. Fitzgerald-Butt, S.M., Bodine, A., Fry, K.M., Ash, J., Zaidi, A.N., Garg, V., Gerhardt, C.A., and McBride, K.L. (2016). Measuring genetic knowledge: a brief survey instrument for adolescents and adults. Clin Genet 89, 235–243. 10.1111/cge.12618.

14. Jallinoja, P., and Aro, A.R. (1999). Knowledge about genes and heredity among Finns. New Genetics and Society 18, 101–110. 10.1080/14636779908656892.

15. Chapman, R., Likhanov, M., Selita, F., and Zakharov, I. (2017). Genetic Literacy And Attitudes Survey (Iglas): International Population-Wide Assessment Instrument 10.15405/epsbs.2017.12.6.

16. Abrams, L.R., Koehly, L.M., Hooker, G.W., Paquin, R.S., Capella, J.N., and McBride, C.M. (2016). Media Exposure and Genetic Literacy Skills to Evaluate Angelina Jolie’s Decision for Prophylactic Mastectomy. Public Health Genomics 19, 282–289. 10.1159/000447944.

17. Ramírez Renta, G.M., Little, I.D., Koehly, L.M., Hilliard, A.J., Foor, K.L., Butts, J., Lundeen, J., and Gunter, C. (2025). Interaction of identity and beliefs with genetic literacy. The American Journal of Human Genetics. 10.1016/j.ajhg.2025.11.014.

18. Calder-Dawe, O., Karen, W., and and Carroll, P. (2020). Being the body in question: young people’s accounts of everyday ableism, visibility and disability. Disability & Society 35, 132–155. 10.1080/09687599.2019.1621742.

19. Bottema-Beutel, K., Kapp, S.K., Lester, J.N., Sasson, N.J., and Hand, B.N. (2021). Avoiding Ableist Language: Suggestions for Autism Researchers. Autism Adulthood 3, 18–29. 10.1089/aut.2020.0014.

20. U.S. Census Bureau Explore Census data.

21. Haga, S.B., Barry, W.T., Mills, R., Ginsburg, G.S., Svetkey, L., Sullivan, J., and Willard, H.F. (2013). Public knowledge of and attitudes toward genetics and genetic testing. Genet Test Mol Biomarkers 17, 327–335. 10.1089/gtmb.2012.0350.

22. Reed, G.F., Lynn, F., and Meade, B.D. (2002). Use of coefficient of variation in assessing variability of quantitative assays. Clin Diagn Lab Immunol 9, 1235–1239. 10.1128/cdli.9.6.1235-1239.2002.

23. Hoang, N., Cytrynbaum, C., and Scherer, S.W. (2018). Communicating complex genomic information: A counselling approach derived from research experience with Autism Spectrum Disorder. Patient Education and Counseling 101, 352–361. 10.1016/j.pec.2017.07.029.

24. R Core Team (2024). R: A language and environment for statistical computing (R Foundation for Statistical Computing).

25. Wickham, H., Hester, J., and Bryan, J. (2024). readr: Read Rectangular Text Data.

26. Wickham, H., François, R., Henry, L., Müller, K., and Vaughan, D. (2023). dplyr: A Grammar of Data Manipulation.

27. Wickham, H., Averick, M., Bryan, J., Chang, W., McGowan, L.D., François, R., Grolemund, G., Hayes, A., Henry, L., Hester, J., et al. (2019). Welcome to the tidyverse. Journal of Open Source Software 4, 1686. 10.21105/joss.01686.

28. Wickham, H. (2007). Reshaping Data with the reshape Package. Journal of Statistical Software 21, 1–20.

29. Fox, J., and Weisberg, S. (2019). An R Companion to Applied Regression, Third Edition (Sage).

30. Hothorn, T., Bretz, F., and Westfall, P. (2008). Simultaneous Inference in General Parametric Models. Biometrical Journal 50, 346–363.

31. Long, J.A. (2022). jtools: Analysis and Presentation of Social Scientific Data.

32. Rosseel, Y. (2012). lavaan: An R Package for Structural Equation Modeling. Journal of Statistical Software 48, 1–36. 10.18637/jss.v048.i02.

33. Epskamp, S., Cramer, A.O.J., Waldorp, L.J., Schmittmann, V.D., and Borsboom, D. (2012). qgraph: Network Visualizations of Relationships in Psychometric Data. Journal of Statistical Software 48, 1–18.

34. Venables, W.N., and Ripley, B.D. (2002). Modern Applied Statistics with S, Fourth Edition (Springer).

35. Wickham, H. (2016). ggplot2: Elegant Graphics for Data Analysis (Springer-Verlag).

36. Kassambara, A. (2023). ggpubr: ‘ggplot2’ Based Publication Ready Plots.

37. Ooms, J. (2025). writexl: Export Data Frames to Excel ‘xlsx’ Format.

38. Robinson, D., Hayes, A., Couch, S. (2024). broom: Convert Statistical Objects into Tidy Tibbles.

39. Howard, M.C. (2016). A Review of Exploratory Factor Analysis Decisions and Overview of Current Practices: What We Are Doing and How Can We Improve? International Journal of Human–Computer Interaction 32, 51–62. 10.1080/10447318.2015.1087664.

40. Li, C.H. (2016). Confirmatory factor analysis with ordinal data: Comparing robust maximum likelihood and diagonally weighted least squares. Behav Res Methods 48, 936–949. 10.3758/s13428-015-0619-7.

41. Hu, L.t., and Bentler, P.M. (1999). Cutoff criteria for fit indexes in covariance structure analysis: Conventional criteria versus new alternatives. Structural Equation Modeling: A Multidisciplinary Journal 6, 1–55. 10.1080/10705519909540118.

42. Hair, J.F., Black, W.C., Babin, B.J., and Anderson, R.E. (2013). Multivariate Data Analysis (Pearson Education Limited).

43. Tavakol, M., and Wetzel, A. (2020). Factor Analysis: a means for theory and instrument development in support of construct validity. Int J Med Educ 11, 245–247. 10.5116/ijme.5f96.0f4a.

44. McCrae, R.R., Kurtz, J.E., Yamagata, S., and Terracciano, A. (2011). Internal consistency, retest reliability, and their implications for personality scale validity. Pers Soc Psychol Rev 15, 28–50. 10.1177/1088868310366253.

45. Fonseca, C., Pettitt, J., Woollard, A., Rutherford, A., Bickmore, W., Ferguson-Smith, A., and Hurst, L.D. (2023). People with more extreme attitudes towards science have self-confidence in their understanding of science, even if this is not justified. PLOS Biology 21, e3001915. 10.1371/journal.pbio.3001915.

46. Lackner, S., Francisco, F., Mendonca, C., Mata, A., and Goncalves-Sa, J. (2023). Intermediate levels of scientific knowledge are associated with overconfidence and negative attitudes towards science. Nat Hum Behav 7, 1490–1501. 10.1038/s41562-023-01677-8.

47. Eysenbach, G., and Wyatt, J. (2002). Using the Internet for surveys and health research. J Med Internet Res 4, E13. 10.2196/jmir.4.2.e13.

48. Toscos, T., Drouin, M., Pater, J., Flanagan, M., Pfafman, R., and Mirro, M.J. (2019). Selection biases in technology-based intervention research: patients’ technology use relates to both demographic and health-related inequities. J Am Med Inform Assoc 26, 835–839. 10.1093/jamia/ocz058.

49. Kruger, J., and Dunning, D. (1999). Unskilled and unaware of it: how difficulties in recognizing one’s own incompetence lead to inflated self-assessments. J Pers Soc Psychol 77, 1121–1134. 10.1037//0022-3514.77.6.1121.

