## Supplementary Materials for "Psychometric Validation of the Education and Assessment of Genetic Literacy (EAGL) Measure"

Table S1. Participant Characteristics for Combined Sample (N = 2708), Split by Sample

Table S2: Comparison of Mean EAGL-Short Scores Across Three Survey Runs

Table S3: Complete Evaluation and Assessment of Genetic Literacy (EAGL-long) Instrument

Table S4: Factor Loadings and Communalities (h^2^) for EAGL-long (Exploratory Factor Analysis, EFA)

Table S5: Five Factor EFA Table

Table S6. Four Factor EFA Table

Table S7: Eigenvalues and Variance Explained for Exploratory Factor Analysis of the EAGL-long

**Figure S1: Exploratory Factor Analysis (EFA) results for the EAGL-long, 3, 4, 5, and 9-factor solutions**

Table S8: Descriptive Statistics for Subjective Knowledge Items

Table S9: Frequency distribution for Knowledge Comprehension and Conceptual Knowledge items in CFA Sample (EAGL3, n = 1001)

Table S10: Average Scores Per Subscale for EAGL-long and EAGL-short

Table S11. Adjusted Mean Estimates for Each Subscale in EAGL-short

| Table S1. Participant Characteristics for Combined Sample (N = 2708), Split by Sample | | | | |
| --- | --- | --- | --- | --- |
| Sample | Variable | Category | Frequency | % Total |
| Sample 1 (N = 1005) | Age | 26-39 | 249 | 24.8% |
|  |  | 50-59 | 217 | 21.6% |
|  |  | 40-49 | 179 | 17.8% |
|  |  | 60-69 | 175 | 17.4% |
|  |  | 18-25 | 136 | 13.5% |
|  |  | 70+ | 49 | 4.9% |
|  | Education | Bachelor's degree | 398 | 39.6% |
|  |  | High school diploma or equivalent | 275 | 27.4% |
|  |  | Professional degree | 159 | 15.8% |
|  |  | Associate degree | 138 | 13.7% |
|  |  | Doctorate degree | 24 | 2.4% |
|  |  | Some high school, no diploma | 9 | 0.9% |
|  |  | Kindergarten through 8th grade | 2 | 0.2% |
|  | Connection to Autism | No | 822 | 81.8% |
|  |  | Yes | 183 | 18.2% |
|  | Metro Status | Metro | 896 | 89.15% |
|  |  | Nonmetro | 108 | 10.75 |
|  |  | Invalid Responses | 1 | 0.10% |
| Sample 2 (n = 702) | Variable | Category | Frequency | % Total |
|  | Age | 18-25 | 134 | 19.1 |
|  |  | 26-39 | 282 | 40.2 |
|  |  | 40-49 | 142 | 20.2 |
|  |  | 50-59 | 91 | 13.0 |
|  |  | 60-69 | 43 | 6.1 |
|  |  | 70+ | 10 | 1.4 |
|  | Education | High school diploma or equivalent | 379 | 54.0% |
|  |  | Associate degree | 145 | 20.7% |
|  |  | Bachelor's degree | 95 | 13.5% |
|  |  | Professional degree | 41 | 5.8% |
|  |  | Doctorate degree | 22 | 3.1% |
|  |  | Some high school, no diploma | 18 | 2.6% |
|  |  | Kindergarten through 8th grade | 2 | 0.3% |
|  | Connection to Autism | No | 523 | 74.5% |
|  |  | Yes | 179 | 25.5% |
|  | Metro Status | Metro | 598 | 85.19% |
|  |  | Nonmetro | 102 | 14.53% |
|  |  | Invalid Responses | 2 | 0.28% |
| Sample 3 (n = 1001) | Variable | Category | Frequency | % Total |
|  | Age | 26-39 | 431 | 43.0% |
|  |  | 40-49 | 217 | 21.7% |
|  |  | 18-25 | 152 | 15.2% |
|  |  | 50-59 | 128 | 12.8% |
|  |  | 60-69 | 54 | 5.4% |
|  |  | 70+ | 19 | 1.9% |
|  | Education | High school diploma or equivalent | 441 | 44.0% |
|  |  | Bachelor's degree | 292 | 29.2% |
|  |  | Associate degree | 124 | 12.4% |
|  |  | Professional degree | 111 | 11.1% |
|  |  | Doctorate degree | 18 | 1.8% |
|  |  | Some high school, no diploma | 14 | 1.4% |
|  |  | Kindergarten through 8th grade | 1 | 0.1% |
|  | Connection to Autism | No | 791 | 79.0% |
|  |  | Yes | 210 | 21.0% |
|  | Metro Status | Metro | 880 | 87.91% |
|  |  | Nonmetro | 120 | 11.99% |
|  |  | Invalid Responses | 1 | 0.10% |

Table S1. The table presents demographic characteristics across three EAGL samples comprising a total of 2,708 participants (Sample 1: N = 1,005; Sample 2: N = 702; Sample 3: N = 1,001). Age categories are presented in years and grouped into six ranges from 18-25 to 70+. Education levels are categorized from elementary education through doctoral degrees. "Connection to Autism" indicates whether participants reported having a personal connection to autism, either through themselves or through family members. Metro/non-metro distinctions were made via Federal Information Processing System (FIPS) Codes for States and Counties and Rural-Urban Continuum Codes (RUCC). A RUCC of 1-3 delineates a metropolitan area, with 4-9 delineating a nonmetropolitan county. The specific numbers indicate the level of adjacency to a metropolitan area, considering population of county and proximity to metro counties. Four participants had to be removed from analysis as they provided invalid 5-digit zip codes. Frequencies represent the number of participants in each category, and percentages are calculated based on the total sample size for each respective sample.

| Table S2: Comparison of Mean EAGL-Short Scores Across Three Survey Runs | | | | | |
| --- | --- | --- | --- | --- | --- |
| Knowledge Domain | EAGL1  Mean (SD) | EAGL2  Mean (SD) | EAGL3  Mean (SD) | Overall Mean (SD) | Coefficient of Variance (CV) (%) |
| Subjective Knowledge | 5.57 (0.98) | 5.59 (0.97) | 5.56 (0.98) | 5.57 (0.97) | 0.29 |
| Applied Knowledge | 7.05 (1.08) | 6.89 (1.13) | 6.93 (1.16) | 6.96 (1.12) | 1.19 |
| Situational Knowledge | 5.56 (1.12) | 5.53 (1.20) | 5.60 (1.19) | 5.56 (1.15) | 0.57 |
| Knowledge Comprehension | 5.47 (0.79) | 5.44 (0.82) | 5.33 (1.08) | 5.41 (0.92) | 1.35 |
| Objective Knowledge | 14.57 (1.81) | 14.30 (1.91) | 14.55 (1.88) | 14.50 (1.86) | 1.04 |

Table S2. The table presents the comparison of mean EAGL-short scores across the three survey runs. The mean score and standard deviation for each subscale of the EAGL-long are presented, along with the coefficient of variance.

**Table S3: Complete Evaluation and Assessment of Genetic Literacy (EAGL-long) Instrument**

First, we would like to see how familiar you are with words related to genetics.

For each word below, please rate how familiar you are with the word. For example, marking “Completely familiar” on the scale reflects that you are entirely familiar with the word, while marking “Not at all familiar” on the scale means that you are not familiar with the word in any capacity. Please select the answer that best reflects your view. Following that rating, you will then be presented with a fill-in-the-blank style question that utilizes a word related to genetics. Please do your best to select the appropriate response for the question, and please do not use the Internet or seek outside help in answering the questions.

| Not at all familiar |  |  | Neither familiar nor unfamiliar |  |  | Completely familiar |
| --- | --- | --- | --- | --- | --- | --- |
| 1 | 2 | 3 | 4 | 5 | 6 | 7 |

| **A1_1.** Genetic  Genetics is the study of how living things receive common traits from previous ____________. | a. generations **(C)** b. experiences c. examinations d. d. achievements |
| --- | --- |
| **A1_2.** Chromosome  A chromosome contains ______________ material. | a. genetic **(C)** b. digestive c. cellular d. d. brain |
| **A1_3.** Susceptibility    Susceptibility to a disease means you _________________ get the disease. | a. eventually will b. definitely will c. possibly will (C) d. d. never will |
| **A1_4** Mutation    A DNA mutation is ______________. | a. a type of cell b. a type of virus c. a change in your DNA sequence (C) d. a measurement of DNA sequence |
| **A1_5.** Variation    Having a variation in the genetic code is __________________. | a. always harmful b. can be harmful or benign (C) c. always benign d. extremely rare |
| **A1_6.** Genome    Your genome is _______________________. | a. where you make new DNA b. all of the genes within one chromosome c. your entire and complete set of DNA (C) d. where cellular respiration occurs |
| **A1_7.** Heredity    Heredity is the transfer of characteristics from ____________________________. | a. the environment to the person b. the sick to the healthy c. the biological parent to their child (C) d. the teacher to their student |
| **A1_8.** Sporadic  If someone is diagnosed with breast cancer without _____________ it is considered sporadic. | a. symptoms b. a tumor c. those around them knowing d. a genetically increased likelihood (C) |
| **A2_1.** Please select the number 2 (two) from the list of numbers below: | a. 1 b. 4 c. 6 d. 2 (C) |

| **B1_1.**  Imagine that we rolled a fair, six-sided die 1,000 times. Out of 1,000 rolls, how many times do you think the die would come up even (2, 4, or 6)? | a. 500 **(C)** b. 250 c. 750 d. 800 |
| --- | --- |
| **B1_2.**  In the BIG BUCKS LOTTERY, the chances of winning a $10.00 prize is 1%. What is your best guess about how many people would win a $10.00 prize if 1,000 people each buy a single ticket to BIG BUCKS? | a. 100 people out of 1000 b. 50 people out of 1000 c. 10 people out of 1000 **(C)** d. 1 person out of 1000 |
| **B1_3.** Which of the following numbers represents the biggest risk of getting a disease? | a. 1 in 100 b. 1 in 10 **(C)** c. 1 in 1000 |

| **C1_1.** You are with your biological aunt, cousin, and sibling. You are asked to line them up by the amount of DNA, or genetic material, you share. You order them by most amount of DNA shared to least. That order is: | a. Aunt, cousin, sibling b. Sibling, aunt, cousin **(C)** c. Sibling, cousin, aunt d. Cousin, sibling, aunt |
| --- | --- |
| **C1_2.** If both of my biological parents have brown eyes, what eye color could I have? | a. Only brown eyes b. Brown or blue eyes c. Brown, blue, or green eyes **(C)** d. Only green eyes |
| **C1_3.** Certain genetic variants in the *BRCA* genes increase someone’s risk for developing breast cancer. What is the most common purpose for obtaining *BRCA* genetic testing? | a. To determine if someone already has developed breast cancer b. To determine if someone will never develop breast cancer c. To determine if someone will one day develop breast cancer due to their environment d. To determine if someone is genetically predisposed to developing breast cancer **(C)** |
| **C1_4.** If you have a genetic disease that was cured in your own red blood cells using gene therapy (replacing defective or missing genes with functional ones) as an adult, can you still pass on the disease to your biological children? | a. Yes, they could still get the disease **(C)** b. No, they could not get the disease |
| **C1_5.** Cloning results in two organisms that are | a. genetically similar b. genetically identical **(C)** c. look alike but have different DNA d. adults |
| **C1_6**. The term DNA stands for | a. Dynamic nitrogenous assembly b. Double-nucleotide association c. Deoxyribonucleic acid **(C)** d. Downward nebula artifact |
| **C1_7**. When scientists say that a trait is “sex-linked,” they mean that the genes related to the trait are | a. On the X or Y chromosome **(C)** b. On all of the chromosomes c. Only present in men d. Only present in women |
| **C2_1.** Please select the third option from the list: | a. a b. b c. c **(C)** d. d |

| **D1_1.** As far as you know, do you or does anyone in your immediate family have a genetic condition? | Yes …......................................................1  No ….......................................................2 |
| --- | --- |
| **D1_2.** As far as you know, have you or an immediate family member (e.g. parent, sibling, child) received any form of genetic testing? | Yes …......................................................1  No ….......................................................2 |

[**INFOGRAPHIC ABOUT GENETIC AND ENVIRONMENTAL CONTRIBUTORS TO AUTISM DIAGNOSIS HERE]**

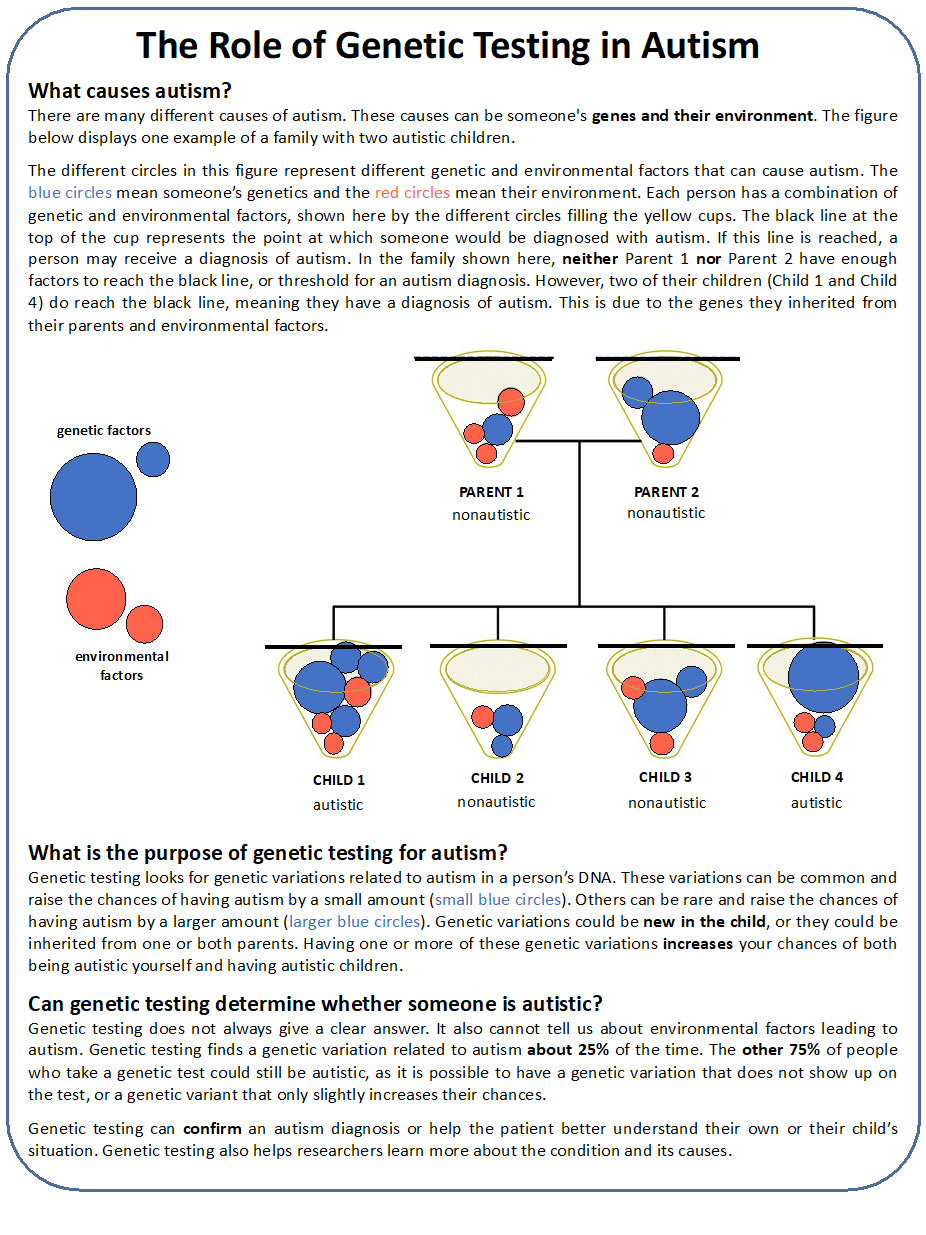

| **[CHECKBOXES]**  **E1.** What is the purpose of genetic testing for autism? | Genetic testing always provides a clear answer to whether or not someone is autistic ………...................................................................................1  Genetic testing can tell you both the genetic and environmental causes of autism …...........................................................................................2  Genetic testing analyzes someone’s DNA to find genetic variations related to autism …................................................................................................................3 **(C)** |
| --- | --- |
| E2. Please select the phrase that best completes the following statement: Genetic variations that could increase a person’s chance of being autistic… | Can be present in some siblings and not others….........................................1 **(C)**  Are always the exact same between siblings…...................................................2 |
| **[NUMERICAL TEXT BOX: RANGE 0-100]**  **E3.** What percentage of individuals who have genetic testing for autism are found to have a variation related to autism? | Enter a number from 0–100: 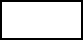% **(25)** |
| [NUMERICAL TEXT BOX: RANGE 0-100]  E4. What percentage of individuals who have genetic testing for autism will receive results with no genetic variations related to autism? | Enter a number from 0–100: 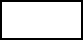% **(75)** |
| **E5.** If neither of the biological parents are autistic, it is impossible for their child to be autistic. | True.............................................................1 False............................................................2 **(C)** |
| **E6.** It is possible for a child to have a genetic variation that raises their chances of being autistic which neither of their biological parents have. | True.............................................................1 **(C)** False............................................................2 |
| E7. Reaching the threshold for an autism diagnosis is due to a variable amount of environmental and genetic factors that vary per individual. | True.............................................................1 **(C)** False............................................................2 |

**[TRUE/FALSE CHOICES]**

| **F1_1.** One can see a gene with the naked eye. | F |
| --- | --- |
| **F1_2.** Healthy parents can have a child with a hereditary condition. | T |
| **F1_3.** The onset of certain diseases is due to genes, environment, and lifestyle. | T |
| **F1_4.** A gene is a disease. | F |
| **F1_5.** The carrier of a gene linked to a disease may not have the disease themselves. | T |
| **F1_6.** All serious diseases are hereditary. | F |
| **F1_7.** Genes control hereditary characteristics. | T |
| **F1_8.** Genes are inside cells. | T |
| **F1_9.** The child of a carrier for a genetic condition is always also a carrier for the same condition. | F |
| **F1_10.** A gene is a piece of DNA. | T |
| **F1_11.** A gene is a cell. | F |
| **F1_12.** A gene is a part of a chromosome. | T |
| **F1_13.** Different genes are expressed in different body parts. | T |
| **F1_14.** Genes are bigger than chromosomes. | F |
| **F1_15.** The genome can be changed by human intervention. | T |
| **F1_16.** It is currently estimated that a person has about 20,000 genes. | T |
| **F1_17.** Environmental factors, such as UV radiation, can change our DNA sequence. | T |

| **G1_1.** What is the most important idea or fact about genetics that you feel you need to know? | **[TEXTBOX RESPONSE]** |
| --- | --- |

| **K1.** How old are you? | < 18 .....................................................................................1  18 - 25 .................................................................................2  26 - 39 .................................................................................3  40 - 49..................................................................................4  50 - 59..................................................................................5  60 - 69 .................................................................................6  70 +......................................................................................7 |
| --- | --- |
| **K2.** What is the highest level of education you have completed? | No schooling completed…...................................................1  Kindergarten through 8^th^ grade.............................................2  Some high school, no diploma .............................................3  High school diploma or equivalent ......................................4  Associate degree....................................................................5  Bachelor’s degree..................................................................6  Professional degree...............................................................7  Doctorate degree...................................................................8 |
| **K3.** Are you or is anyone in your immediate family autistic? | Yes .........................................................1  No ..........................................................2 |
| **K4.** What is the ZIP code of your primary residence? Enter a 5-digit ZIP Code. | **[OPEN ENDED BOX]** |
| **K5.** Did you answer this survey honestly and to the best of your ability? | Yes .........................................................1  No ..........................................................2 |

END

Sections:

A1-8 Likert-scale questions measure subjective knowledge

A1-8 multiple choice questions measure applied knowledge

B1-3 questions measure objective numeracy

C1-7 questions measure situational knowledge

E1-7 questions measure knowledge comprehension

F1-17 questions measure objective knowledge

G1 is an optional qualitative comment area

K1-K5 are demographics questions

Questions contained in the EAGL-short measure are shown in Figure 1 of the paper.

| Table S4. Factor Loadings and Communalities (h^2^) for EAGL-long (Exploratory Factor Analysis) | | | | | | | | | | |
| --- | --- | --- | --- | --- | --- | --- | --- | --- | --- | --- |
| Item | Factor1 | Factor2 | Factor3 | Factor4 | Factor5 | Factor6 | Factor7 | Factor8 | Factor9 | h^2^ |
| GENE | 0.886* | 0.011 | -0.03 | -0.240* | -0.02 | -0.022 | 0.017 | -0.007 | 0.054 | 0.848 |
| CHRO | 0.833* | -0.014 | -0.029 | -0.204* | 0 | 0.137* | 0.034 | -0.006 | 0.023 | 0.757 |
| SUSC | 0.681* | -0.01 | 0.110* | 0.124* | 0.002 | -0.021 | -0.039 | 0.054 | 0.029 | 0.497 |
| MUTA | 0.811* | 0.037 | 0.065 | 0.017 | 0.028 | 0 | -0.032 | 0.018 | -0.123* | 0.681 |
| VARI | 0.562* | -0.065 | -0.034 | 0.309* | 0.059 | 0.041 | -0.004 | 0.055 | -0.011 | 0.425 |
| GENO | 0.407* | 0.035 | -0.045 | 0.132 | -0.009 | 0.648* | 0.017 | 0.002 | -0.009 | 0.607 |
| HERE | 0.709* | 0.013 | 0.120* | 0.001 | 0.011 | -0.027 | -0.026 | -0.085 | -0.08 | 0.532 |
| SPOR | 0.373* | 0.002 | -0.004 | 0.571* | -0.027 | 0 | 0.016 | -0.042 | 0.027 | 0.469 |
| E1 | 0.06 | 0.316* | 0.448* | 0 | 0.124 | -0.027 | -0.057 | -0.013 | 0.049 | 0.326 |
| E2 | -0.002 | 0.038 | 0.582* | -0.031 | 0.002 | -0.156 | 0.055 | 0 | 0.258 | 0.435 |
| E3 | -0.008 | 1.017* | -0.008 | 0.015 | 0.016 | -0.038 | 0.063 | -0.004 | -0.01 | 1.040 |
| E4 | 0.011 | 0.973* | 0.025 | -0.027 | -0.049 | 0.015 | -0.026 | 0.032 | -0.005 | 0.953 |
| E5 | -0.024 | 0.283* | -0.069 | -0.022 | 0.247* | 0.039 | -0.041 | 0.025 | 0.038 | 0.152 |
| E6 | -0.007 | 0.019 | 0.045 | -0.065 | -0.328 | -0.039 | -0.066 | 0.393* | 0.2 | 0.315 |
| F8 | -0.005 | -0.024 | 0.490* | 0.024 | -0.212 | 0.181 | 0.263* | 0.026 | -0.161 | 0.415 |
| F9 | 0.046 | 0.195* | 0.058 | 0.064 | 0.044 | 0.084 | 0.041 | 0.044 | 0.244* | 0.120 |
| F11 | 0.024 | 0.01 | 0.066 | 0.027 | 0.645* | -0.01 | -0.031 | 0.096 | -0.012 | 0.432 |
| F12 | -0.025 | -0.014 | -0.008 | -0.012 | 0.052 | -0.017 | 0.776* | -0.015 | 0.002 | 0.606 |
| F13 | 0.066 | -0.123 | -0.072 | 0 | 0.008 | -0.033 | 0.019 | 0.346* | 0.058 | 0.149 |
| F14 | 0.014 | 0.03 | 0.04 | -0.013 | 0.368* | 0.007 | 0.512* | 0 | 0.037 | 0.402 |
| F15 | 0.034 | 0.01 | -0.028 | 0.027 | -0.045 | 0.162* | 0.053 | 0.512* | -0.024 | 0.297 |
| F16 | 0.058 | -0.150 | -0.008 | -0.011 | -0.093 | -0.022 | 0.143* | 0.093 | 0.026 | 0.065 |
| F17 | -0.008 | 0.037 | 0.006 | 0.01 | 0.065 | 0.014 | 0.004 | 0.721* | -0.361* | 0.656 |
| A5 | -0.03 | 0.066 | 0.174 | 0.08 | 0.079 | -0.014 | -0.045 | 0.300* | 0.101 | 0.151 |
| A6 | 0.001 | -0.02 | 0.361* | -0.038 | 0.067 | 0.561* | -0.219 | -0.011 | 0.023 | 0.500 |
| A8 | 0.002 | 0.079 | 0.294* | 0.171* | 0.037 | 0.078 | 0.041 | 0.092 | 0.072 | 0.145 |
| C2 | 0.002 | -0.002 | 0.087 | 0.035 | 0.038 | 0.186 | -0.002 | 0.026 | 0.503* | 0.299 |
| C4 | -0.018 | 0.021 | -0.029 | 0.078 | -0.001 | -0.064 | -0.07 | -0.042 | 0.224* | 0.069 |
| C5 | 0.046 | 0.05 | 0.022 | 0.028 | 0.399* | 0.071 | 0.06 | 0.022 | 0.022 | 0.175 |
| C6 | -0.01 | -0.067 | 0.041 | -0.047 | 0.343* | 0.281* | 0.041 | 0.247* | -0.027 | 0.269 |

Table S4. This table presents the results of an exploratory factor analysis (EFA) for the EAGL-long instrument, displaying factor loadings and communalities across nine extracted factors (F1-F9). Each row represents a scale question. Factor loadings indicate the correlation between each item and the underlying factors, with values closer to ±1.0 representing stronger relationships. Asterisks (*) denote statistically significant factor loadings. The communality (h²) represents the proportion of each item's variance explained by the factor solution, with values ranging from 0 to 1, where higher values indicate that more of the item's variance is accounted for by the extracted factors. Survey questions F1-F7 were excluded from this 9-factor solution due to excessively high accuracy rates, suggesting potential issues with model fit or interpretability.

| Table S5. Five Factor EFA Table | | | | | |
| --- | --- | --- | --- | --- | --- |
| Variable | Factor1 | Factor2 | Factor3 | Factor4 | Factor5 |
| GENE | 0.891* | 0.032* | -0.097* | -0.196* | 0.007 |
| CHRO | 0.857* | -0.058* | 0.052* | -0.173* | 0.02 |
| SUSC | 0.666* | 0.100* | 0.038 | 0.142* | -0.035 |
| MUTA | 0.812* | 0.049* | 0.018 | 0.013 | -0.01 |
| VARI | 0.546* | -0.059* | 0.102* | 0.305* | -0.006 |
| GENO | 0.455* | -0.194* | 0.439* | 0.149* | 0.007 |
| HERE | 0.711* | 0.056* | -0.069* | 0.019 | -0.012 |
| SPOR | 0.344* | 0.011 | -0.037* | 0.586* | 0.015 |
| E1N | 0.058 | 0.578* | 0.187* | 0.016 | -0.028 |
| E2N | -0.04 | 0.493* | 0.088 | 0.028 | 0.044 |
| E3N | 0 | 0.976* | -0.054* | -0.012 | 0.045 |
| E4N | 0.019 | 0.925* | -0.024 | -0.044 | -0.028 |
| E5N | -0.011 | 0.299* | 0.181* | -0.032 | -0.038 |
| E6N | -0.048 | 0.138 | 0.041 | -0.058 | -0.053 |
| F8N | -0.005 | 0.032 | 0.205* | 0.025 | 0.236* |
| F9N | 0.043 | 0.306* | 0.141* | 0.098* | 0.021 |
| F11N | 0.038 | 0.226* | 0.492* | -0.002 | -0.009 |
| F12N | -0.012 | -0.032 | -0.014 | 0.011 | 0.906* |
| F13N | 0.033 | -0.043 | 0.216* | -0.026 | 0.011 |
| F14N | 0.021 | 0.125* | 0.290* | -0.011 | 0.431* |
| F15N | 0.004 | 0.002 | 0.479* | -0.018 | 0.036 |
| F16N | 0.045 | -0.134* | -0.01 | -0.008 | 0.136* |
| F17N | -0.045 | 0.033 | 0.549* | -0.086 | 0.035 |
| A5N | -0.063 | 0.284* | 0.331* | 0.058 | -0.051 |
| A6N | 0.054 | 0.006 | 0.551* | 0.017 | -0.157* |
| A8N | -0.013 | 0.263* | 0.257* | 0.181* | 0.049 |
| C2N | 0.013 | 0.205* | 0.192* | 0.108* | -0.022 |
| C4N | -0.026 | 0.133* | -0.108 | 0.112* | -0.067 |
| C5N | 0.058 | 0.150* | 0.348* | 0.021 | 0.062 |
| C6N | -0.002 | -0.039 | 0.688* | -0.08 | 0.02 |

Table S5. Exploratory factor analysis showing factor loadings for EAGL-short scale items across five factors. Asterisks (*) indicate statistically significant loadings. Variables represent different dimensions of genetic knowledge and attitudes, with loadings ranging from -0.196 to 0.976.

| Table S6. Four Factor EFA Table | | | | |
| --- | --- | --- | --- | --- |
| Variable | Factor1 | Factor2 | Factor3 | Factor4 |
| GENE | 0.833* | 0.044* | -0.166* | 0.073* |
| CHRO | 0.806* | -0.043* | -0.021 | 0.092* |
| SUSC | 0.699* | 0.086* | 0.100* | -0.063* |
| MUTA | 0.821* | 0.041* | 0.025 | -0.004 |
| VARI | 0.609* | -0.080* | 0.214* | -0.065* |
| GENO | 0.488* | -0.216* | 0.488* | 0.012 |
| HERE | 0.722* | 0.053* | -0.062* | -0.008 |
| SPOR | 0.456* | -0.02 | 0.167* | -0.096* |
| E1N | 0.054 | 0.572* | 0.213* | -0.039 |
| E2N | -0.038 | 0.482* | 0.114 | 0.03 |
| E3N | -0.004 | 0.974* | -0.044* | 0.049 |
| E4N | 0.007 | 0.930* | -0.027 | -0.01 |
| E5N | -0.022 | 0.303* | 0.170* | -0.013 |
| E6N | -0.066 | 0.144 | 0.029 | -0.051 |
| F8N | 0.003 | 0.012 | 0.199* | 0.248* |
| F9N | 0.061 | 0.289* | 0.194* | -0.013 |
| F11N | 0.034 | 0.224* | 0.474* | 0.054 |
| F12N | -0.014 | -0.086 | 0 | 0.700* |
| F13N | 0.026 | -0.039 | 0.190* | 0.05 |
| F14N | 0.025 | 0.104* | 0.239* | 0.519* |
| F15N | -0.002 | 0.002 | 0.444* | 0.106 |
| F16N | 0.045 | -0.141* | -0.022 | 0.136* |
| F17N | -0.067 | 0.043 | 0.479* | 0.135* |
| A5N | -0.054 | 0.280* | 0.355* | -0.038 |
| A6N | 0.048 | 0.005 | 0.560* | -0.123* |
| A8N | 0.025 | 0.235* | 0.337* | 0.003 |
| C2N | 0.032 | 0.188* | 0.251* | -0.058 |
| C4N | -0.004 | 0.122* | -0.036 | -0.139* |
| C5N | 0.059 | 0.140* | 0.347* | 0.09 |
| C6N | -0.021 | -0.026 | 0.607* | 0.137* |

Table S6. Exploratory factor analysis showing factor loadings for EAGL-short scale items across four factors. Asterisks (*) indicate statistically significant loadings. Variables represent different dimensions of genetic knowledge and attitudes, with loadings ranging from -0.166 to 0.974.

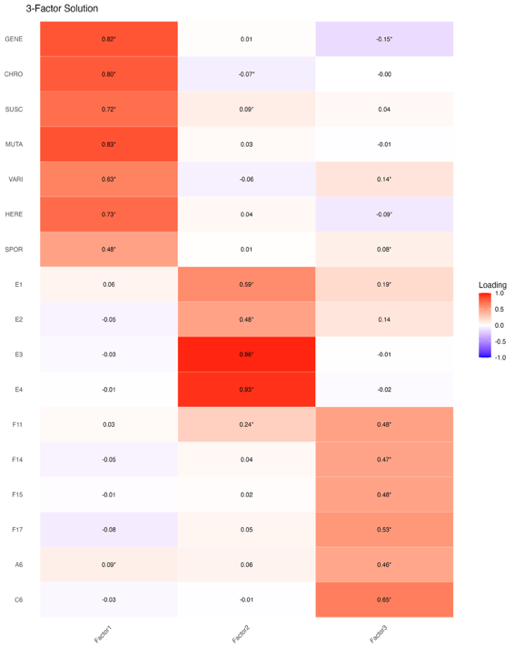

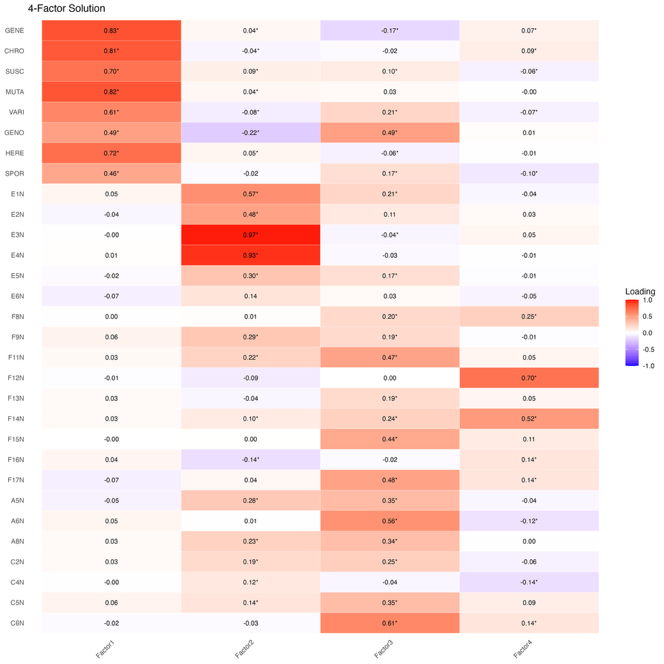

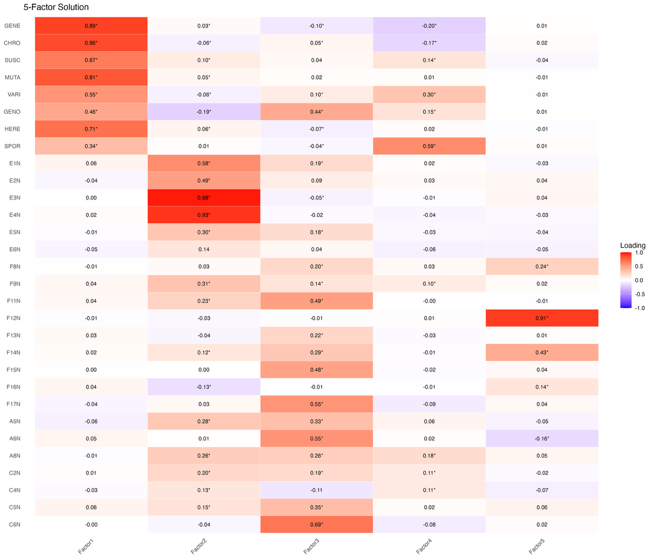

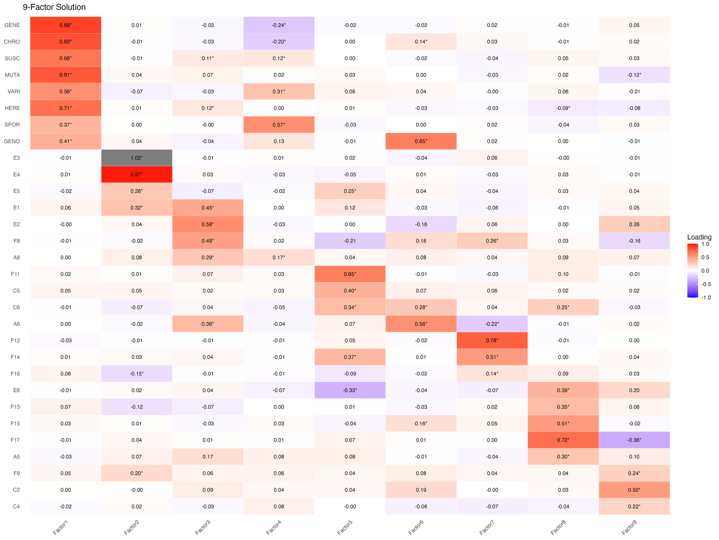

a

c

b

d

**Figure S1. Exploratory Factor Analysis (EFA) results for the EAGL-long, 3, 4, 5, and 9-factor solutions.** The heatmap displays factor loadings, with color intensity representing the magnitude of loadings (red for positive, blue for negative), and asterisks (*) indicating statistically significant loadings. Questions are grouped to highlight primary factor associations, with some items loading on multiple factors significantly, or loading on none. **(a)** 3-factor solution. **(b)** 4-factor solution. **(c)** 5-factor solution. **(d)** 9-factor solution contains nuanced splicing of clusters which illustrate multiple subject domains being tested. Loading values represent the strength of association between each item and the factor.

| Table S7. Eigenvalues and Variance Explained for Exploratory Factor Analysis of the EAGL-long | | | |
| --- | --- | --- | --- |
| Factor | Eigenvalue | % of Variance | Cumulative % |
| Factor 1 | 3.744 | 12.48 | 12.50 |
| Factor 2 | 2.268 | 7.56 | 20.0 |
| Factor 3 | 1.096 | 3.65 | 23.70 |
| Factor 4 | 0.615 | 2.05 | 25.70 |
| Factor 5 | 1.099 | 3.66 | 29.4 |
| Factor 6 | 0.984 | 3.28 | 32.7 |
| Factor 7 | 1.044 | 3.48 | 36.2 |
| Factor 8 | 1.257 | 4.19 | 40.4 |
| Factor 9 | 0.680 | 2.27 | 42.60 |

Table S7. This table presents the eigenvalues and variance explained for each factor identified through Exploratory Factor Analysis of the EAGL-long measure. Each row represents a factor, ordered from highest to lowest eigenvalue. The first column presents the eigenvalues, or how much variance each factor explains. Those above the Kaiser criterion (eigenvalues > 1) are considered to be significantly meaningful. The second column presents this another way, through the percentage of total variance in the data that each factor accounts for individually. The third column, cumulative percentage, shows the total variance explained when you include each factor with the previous factors listed. All nine factors together explain 42.60% of the total variance in the data.

| Table S8. Descriptive Statistics for Subjective Knowledge Items | | | | |
| --- | --- | --- | --- | --- |
|  | Mean | SD | Skewness | Kurtosis |
| Genetic | 6.09 | 0.99 | -1.13 | 1.59 |
| Chromosome | 5.76 | 1.12 | -0.86 | 0.99 |
| Susceptibility | 5.73 | 1.29 | -1.25 | 1.87 |
| Mutation | 5.89 | 1.15 | -1.24 | 2.07 |
| Variation | 5.38 | 1.46 | -0.94 | 0.61 |
| Heredity | 6.1 | 1.04 | -1.45 | 3.13 |
| Sporadic | 4.92 | 1.68 | -0.66 | -0.31 |
| Overall | 5.69 | 0.95 | -0.69 | 0.48 |

Table S8. This table presents the descriptive statistics for the subjective knowledge items in the EAGL-short. Items were presented on a Likert scale, ranging from not at all familiar (0) to completely familiar (7). Mean scores, standard deviation, skewness, and kurtosis are presented.

| Table S9. Frequency distribution for Knowledge Comprehension and Conceptual Knowledge items in CFA Sample (EAGL3, n = 1001) | | | | | | |
| --- | --- | --- | --- | --- | --- | --- |
| Factor | Item | 1 (correct) | % | 0 (incorrect) | % | n |
| Knowledge Comprehension | D1 (E1) | 912 | 91.11 | 89 | 8.99 | 1001 |
|  | D2 (E2) | 979 | 97.80 | 22 | 2.20 | 1001 |
|  | D3 (E3) | 868 | 86.71 | 133 | 13.29 | 1001 |
|  | D4 (E4) | 835 | 83.42 | 166 | 16.58 | 1001 |
| Conceptual Knowledge | E1_1 (F11) | 735 | 73.43 | 266 | 26.57 | 1001 |
|  | E1_2 (F14) | 760 | 75.92 | 241 | 24.08 | 1001 |
|  | E1_3 (F15) | 718 | 71.73 | 283 | 28.27 | 1001 |
|  | E1_4 (F17) | 807 | 80.82 | 194 | 19.38 | 1001 |
|  | E2_1 (A6) | 745 | 74.43 | 256 | 25.57 | 1001 |
|  | E2_2 (C6) | 758 | 75.72 | 243 | 24.28 | 1001 |

Table S9. This table presents the frequency distributions for knowledge comprehensions and conceptual knowledge items within the sample used for CFA (EAGL3 sample, n = 1001). EAGL1 and EAGL2 were utilized for exploratory factor analysis (EFA) to identify the underlying factor structure, and EAGL3 for CFA to confirm the identified structure. This approach ensures a more rigorous validation of the factor model. Frequencies and percentages show the number and proportion of participants who answered each item correctly (1) or incorrectly (0). Items D1-D4 measure Knowledge Comprehension, while items E1_1-E2_2 measure Conceptual Knowledge as identified in exploratory factor analysis.

| Table S10. Average Scores Per Subscale for EAGL-long and EAGL-short | | | | |
| --- | --- | --- | --- | --- |
| Instrument | Subscale | Mean | SD | Total Possible Score |
| EAGL-long | Subjective Knowledge | 5.57 | 0.97 | 8 |
|  | Applied Knowledge | 6.96 | 1.12 | 8 |
|  | Situational Knowledge | 5.57 | 1.15 | 7 |
|  | Knowledge Comprehension | 5.41 | 0.91 | 6 |
|  | Objective Knowledge | 14.49 | 1.86 | 17 |
|  | Total Overall Score | 38 | 3.86 | 46 |
| EAGL-short | Subjective Knowledge | 5.71 | 0.95 | 7 |
|  | Knowledge Comprehension | 3.68 | 0.70 | 4 |
|  | Conceptual Knowledge | 4.5 | 1.40 | 6 |
|  | Total Overall Score | 13.89 | 2.01 | 17 |

Table S10. This table presents the average scores per subscale for both the EAGL-long and EAGL-short measures. The mean and standard deviation are presented for each subscale, along with for the overall score within each measure. The total possible scores are also presented.

| Table S11. Adjusted Mean Estimates for Each Subscale in EAGL-short | | | | |
| --- | --- | --- | --- | --- |
| Subjective Knowledge | Variable | Value | Mean | Std. Err. |
|  | Age | 26-39 | 5.870501 | 0.048366 |
|  |  | 40-49 | 5.956 | 0.055027 |
|  |  | 50-59 | 5.886153 | 0.058928 |
|  |  | 60-69 | 5.756377 | 0.069122 |
|  |  | 70+ | 5.971 | 0.113743 |
|  |  | 18-25 | 5.789449 | 0.06097 |
|  | Education | High school diploma or equivalent | 5.725129 | 0.03996 |
|  |  | Associate degree | 5.799588 | 0.054729 |
|  |  | Bachelor’s degree | 5.836788 | 0.046494 |
|  |  | Professional degree | 5.947521 | 0.062135 |
|  |  | Doctorate degree | 5.953307 | 0.121663 |
|  |  | Less than High School | 5.967149 | 0.143305 |
|  | Metro Status | Metro | 5.821216 | 0.040745 |
|  |  | Nonmetro | 5.921945 | 0.062807 |
|  | Connection to Autism | Yes | 5.966647 | 0.056015 |
|  |  | No | 5.776513 | 0.043638 |
| Knowledge Comprehension | Age | 26-39 | 3.643372 | 0.039485 |
|  |  | 40-49 | 3.69233 | 0.043838 |
|  |  | 50-59 | 3.732049 | 0.046419 |
|  |  | 60-69 | 3.760242 | 0.053693 |
|  |  | 70+ | 3.740388 | 0.085899 |
|  |  | 18-25 | 3.65797 | 0.048235 |
|  | Education | High school diploma or equivalent | 3.713923 | 0.031255 |
|  |  | Associate degree | 3.713519 | 0.045621 |
|  |  | Bachelor’s degree | 3.774669 | 0.039513 |
|  |  | Professional degree | 3.552954 | 0.05662 |
|  |  | Doctorate degree | 3.819741 | 0.117567 |
|  |  | Less than High School | 3.651545 | 0.127593 |
|  | Metro Status | Metro | 3.697003 | 0.034415 |
|  |  | Nonmetro | 3.71178 | 0.049059 |
|  | Connection to Autism | Yes | 3.720518 | 0.059353 |
|  |  | No | 3.688265 | 0.034312 |
| Conceptual Knowledge | Age | 26-39 | 4.408883 | 0.087295 |
|  |  | 40-49 | 4.322871 | 0.094327 |
|  |  | 50-59 | 4.345177 | 0.100913 |
|  |  | 60-69 | 4.303135 | 0.112935 |
|  |  | 70+ | 4.442354 | 0.174413 |
|  |  | 18-25 | 4.448759 | 0.102803 |
|  | Education | High school diploma or equivalent | 4.432778 | 0.065618 |
|  |  | Associate degree | 4.440231 | 0.104713 |
|  |  | Bachelor’s degree | 4.656137 | 0.097307 |
|  |  | Professional degree | 4.333828 | 0.156439 |
|  |  | Doctorate degree | 4.630176 | 0.242011 |
|  |  | Less than High School | 3.77803 | 0.333141 |
|  | Metro Status | Metro | 4.501379 | 0.06269 |
|  |  | Nonmetro | 4.255681 | 0.14664 |
|  | Connection to Autism | Yes | 4.38015 | 0.095447 |
|  |  | No | 4.37691 | 0.081367 |

Table S11. Supplementary table 11 presents the adjusted mean estimates for each demographic group across three subscales of the EAGL-short measure: subjective knowledge, knowledge comprehension, and conceptual knowledge. Columns from left to right present the variables of interest, values of interests, adjusted means and standard error. Adjusted mean estimates control for the influence of other demographic variables on the variable of interest. Higher adjusted means indicate higher average scores within each subscale for the respective demographic group.
